# EPHA4 signaling dysregulation links abnormal locomotion and the development of idiopathic scoliosis

**DOI:** 10.1101/2024.03.04.583361

**Authors:** Lianlei Wang, Xinyu Yang, Sen Zhao, Pengfei Zheng, Wen Wen, Kexin Xu, Xi Cheng, Qing Li, Anas M. Khanshour, Yoshinao Koike, Junjun Liu, Xin Fan, Nao Otomo, Zefu Chen, Yaqi Li, Lulu Li, Haibo Xie, Panpan Zhu, Xiaoxin Li, Yuchen Niu, Shengru Wang, Sen Liu, Suomao Yuan, Chikashi Terao, Ziquan Li, Shaoke Chen, Xiuli Zhao, Pengfei Liu, Jennifer E. Posey, Zhihong Wu, Guixing Qiu, DISCO study group (Deciphering Disorders Involving Scoliosis & COmorbidities), Shiro Ikegawa, James R. Lupski, Jonathan J. Rios, Carol A. Wise, Terry Jianguo Zhang, Chengtian Zhao, Nan Wu

## Abstract

Idiopathic scoliosis (IS) is the most common form of spinal deformity with unclear pathogenesis. In this study, we firstly reanalyzed the loci associated with IS, drawing upon previous studies. Subsequently, we mapped these loci to candidate genes using either location-based or function-based strategies. To further substantiate our findings, we verified the enrichment of variants within these candidate genes across several large IS cohorts encompassing Chinese, East Asian, and European populations. Consequently, we identified variants in the *EPHA4* gene as compelling candidates for IS. To confirm their pathogenicity, we generated zebrafish mutants of *epha4a*. Remarkably, the zebrafish *epha4a* mutants exhibited pronounced scoliosis during later stages of development, effectively recapitulating the IS phenotype. We observed that the *epha4a* mutants displayed defects in left-right coordination during locomotion, which arose from disorganized neural activation in these mutants. Our subsequent experiments indicated that the disruption of the central pattern generator (CPG) network, characterized by abnormal axon guidance of spinal cord interneurons, contributed to the disorganization observed in the mutants. Moreover, when knocked down *efnb3b*, the ligand for Epha4a, we observed similar CPG defects and disrupted left-right locomotion. These findings strongly suggested that ephrin B3-Epha4 signaling is vital for the proper functioning of CPGs, and defects in this pathway could lead to scoliosis in zebrafish. Furthermore, we identified two cases of IS in *NGEF*, a downstream molecule in the EPHA4 pathway. Collectively, our data provide compelling evidence that neural patterning impairments and disruptions in CPGs may underlie the pathogenesis of IS.

## Introduction

Idiopathic scoliosis (IS) is the most common form of spinal deformity, affecting about 2.5% of the global population (Hresko, 2013; Luk et al., 2010). IS may have long-term physical and mental health consequences, such as cosmetic deformity, cardiopulmonary impairment, and even disability (Weinstein et al., 2003). All these consequences can severely reduce the quality of life. Early intervention with conservative treatments, such as braces, can control scoliosis progression and reduce the need for surgical intervention (Weinstein et al., 2013). However, IS often remains undiagnosed until malformation is evident, emphasizing the importance of risk-prediction measurements in medical management.

Genetic factors are thought to play a significant role in the development of IS, while only a few genes have been associated with the condition to date (Cheng et al., 2015; Miller, 2007). Several single-nucleotide polymorphisms (SNPs) associated with susceptibility to IS have been identified through genome-wide association studies (GWASs), including SNPs linked to genes such as *LBX1*, *GPR126*, and *BNC2*. Notably, knockdown or overexpression of zebrafish homologs of these IS-associated genes have yielded body axis defects (Guo et al., 2016; Kou et al., 2019; Kou et al., 2013; Ogura et al., 2015; Zhu et al., 2015). In addition to common SNPs, rare variants with larger effect sizes or causing rare Mendelian disease traits also contribute to IS. Rare variants in *FBN1*, *FBN2,* and other extracellular matrix genes are associated with severe IS (Buchan et al., 2014; Haller et al., 2016). Linkage analysis of familial IS suggests that it follows autosomal dominant (AD), X-linked dominant, and multifactorial patterns of inheritance (Miller et al., 2005). Variants in *CHD7* and *AKAP2* are also implicated in the pathogenesis of Mendelian forms of IS (Gao et al., 2007; Liebeskind et al., 2016). Finally, centriolar protein POC5 and planar cell polarity protein Vang-like protein 1 (VANGL1) were also shown to be associated with IS. Notably, all these genes participate in a wide range of biological processes, the mechanisms underlying IS progression still require further investigation. Consequently, there is currently no consensus on the etiology of IS (Tang et al., 2021).

In this study, we developed a novel pipeline that combines SNP-to-gene mapping and rare variant association analysis. Using this approach, we analyzed a large Chinese population with IS and further validated our findings in East Asian and European populations. Through these analyses, we identified *EPHA4* as a novel gene associated with IS. To confirm the pathogenicity of this gene, we used a zebrafish model and found that impaired EPHA4 pathway components and resulting defects in central pattern generators (CPGs) were associated with asymmetric spinal locomotion, which may serve as a contributory factor in the development of IS. Additionally, we searched for candidate genes related to EPHA4 signaling pathways and identified mutations in the *NGEF* among IS patients. Overall, our data suggest that the impairment of the EPHA4 pathway and central pattern generators plays a previously unknown role in the development and progression of IS.

## Results

### Enrichment analysis of rare variants in IS candidate genes

To identify candidate genes associated with IS, we conducted a comprehensive literature review of SNPs that are linked to IS from 14 published GWASs. This led us to obtain forty-one SNPs that showed genome-wide significance (P < 5×10^−8^) as detailed in Table S1. We then employed positional mapping, expression quantitative trait locus (eQTL) mapping, or chromatin interaction mapping to identify 156 candidate genes that could be potentially linked to these SNPs. Further analysis of rare variants for the 156 candidate genes using exome data from 411 Chinese probands and 3,800 unrelated Chinese controls identified *EPHA4* as the only significant gene (P=0.045, OR=4.09) (Table S2). EPHA4 is a member of the Eph family of receptor tyrosine kinases which play a vital role in the development of the nervous system (Kania and Klein, 2016). We identified three rare variants in *EPHA4* from three IS patients, including one splicing-donor variant and two missense variants (Table S2). The flowchart of the entire pipeline is shown in Fig. S1.

### Inheritance pattern and functional analyses of variants in *EPHA4*

We characterized the inheritance pattern of variants in *EPHA4* using either trio sequencing data or Sanger sequencing for the parents. None of the parents being tested had scoliosis based on a clinical screening test. Of the three rare variants in *EPHA4*, two (NM_004438.3: c.1443+1G>C and c.2546G>A [p.Cys849Tyr]) arose *de novo* (Fig. 1A and Table 1). The heterozygous splicing variant (c.1443+1G>C) was identified in a female IS patient (SCO2003P0846) (Fig. 1B). Scoliosis in this patient was diagnosed in her 10s, with a main curve Cobb angle of 60°. The *in vitro* minigene assay showed that the c.1443+1G>C variant introduced a new splicing site, resulting in a 36-bp in-frame deletion in exon 6 (Fig. 1G, 1I, S2A). The *EPHA4* heterozygous missense variant (c.2546G>A, p.Cys849Tyr), identified in a female IS patient (SCO2003P2146) (Fig. 1C), is located in the protein tyrosine kinase domain of EPHA4 protein (Fig. 1A). The onset of scoliosis in this patient was in her 10s with a 70° main curve Cobb angle. Given the critical role of EPHA4 in phosphorylating CDK5, which in turn activates downstream signaling pathways (Fu et al., 2007), a western blot analysis of *EPHA4*-c.2546G>A variant was conducted to investigate the protein expression levels of EPHA4 and CDK5 and the amount of phosphorylated CDK5 (pCDK5) in HEK293T cells transfected with *EPHA4*-mutant or *EPHA4*-WT plasmid. Our results revealed that the missense variant resulted in a decreased phosphorylation level of CDK5 (p=0.015), suggesting a partial loss-of-function (LoF) of *EPHA4* (Fig. 1F).

**Figure 1.**
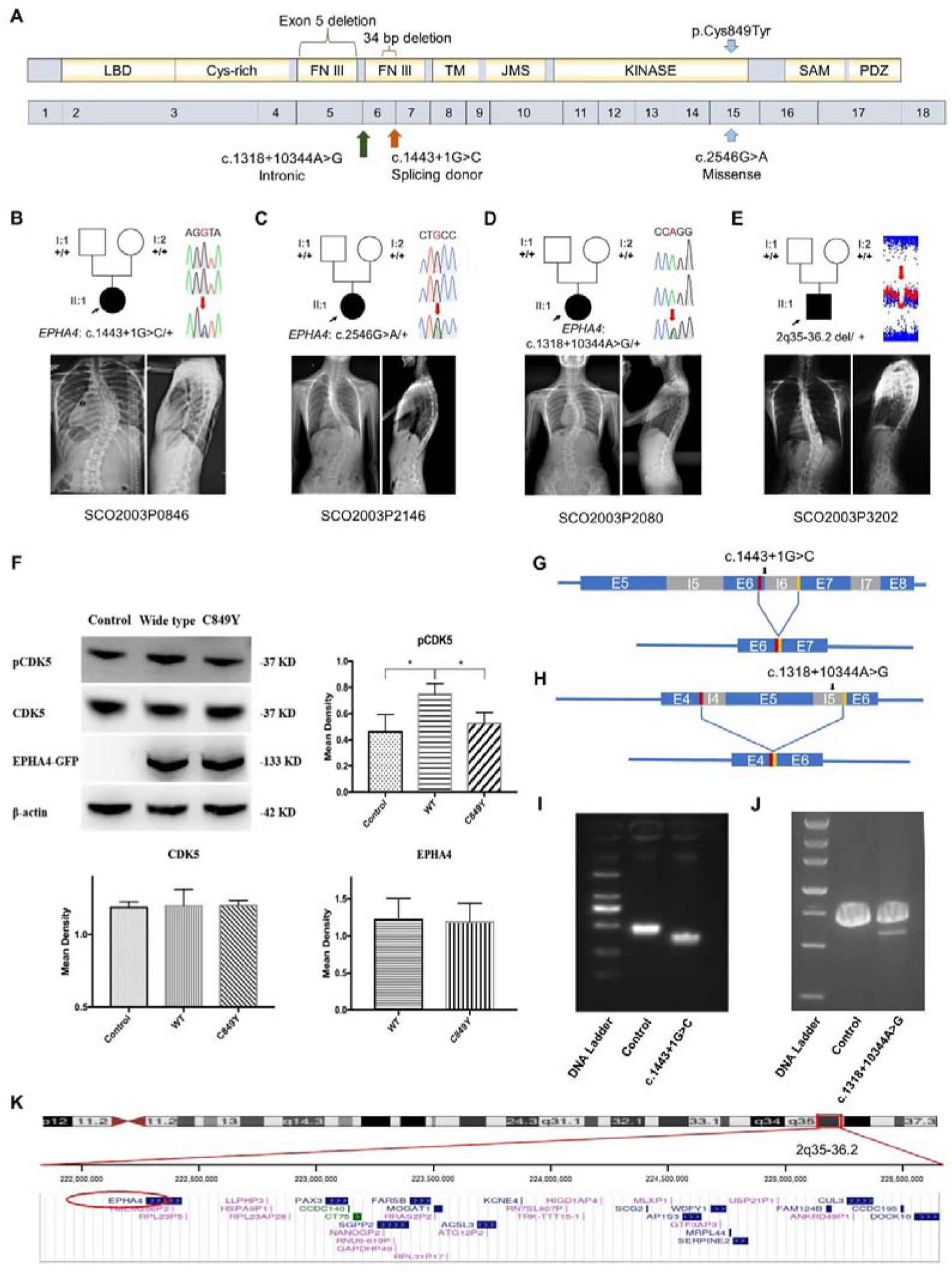
Clinical and genetic information on IS patients and functional effect of *EPHA4* variants. **A:** Locations of three *EPHA4* single nucleotide variants relative to the protein domains (top panel) and exons 1-18 (bottom panel). **B-E:** Pedigrees and spinal radiographs of four probands with dominant gene variants. Sanger sequencing confirmed the variants. The arrows indicate the probands. The term +/+ denotes the wild-type, and cDNA change/+ denotes the heterozygous variant. **F:** Western blot analysis of *EPHA4*-c.2546G>A variant showing the protein expression levels of EPHA4 and CDK5 and the amount of phosphorylated CDK5 (pCDK5) in HEK293T cells transfected with *EPHA4*-mutant or *EPHA4*-WT plasmid. WT: wild type. **G:** Schematic representation of the effect of the *EPHA4*-c.1443+1G>C mutation on the splicing process. This variant induced a new splicing site (red box). The yellow box indicates the splicing acceptor. **H:** Schematic representation of the effect of the *EPHA4*-c.1318+10344A>G mutation on the splicing process. This variant induced a new splicing site (red box). The yellow box indicates the splicing acceptor. **I:** The minigene assay result showed that the c.1443+1G>C variant introduced a new splicing site, resulting in a 36-bp in-frame deletion in exon 6. **J:** The nested PCR showed that the c.1318+10344A>G variant induced exon 5 skipping, resulting in a 339-bp in-frame deletion. **K:** NCBI RefSeq genes included in 2q35-q36.2 from UCSC Genome Browser. *EPHA4* is shown by the red oval.

**Table 1.** Dominantly inherited variants identified in *EPHA4* and *NGEF*.

| Patient ID | Chr | Gene | Ethnicity | Inheritance<br>pattern | cDNA change | AA change | Variant<br>type | ExAC<br>PLI | CADD | GnomAD<br>frequency | In-house<br>frequency |
| --- | --- | --- | --- | --- | --- | --- | --- | --- | --- | --- | --- |
| SCO2003P0846 | 2 | *EPHA4* | Chinese | De novo | c.1443+1G>C | NA | Splice<br>donor | 1 | 21.4 | 0 | 0 |
| SCO2003P2146 | 2 | *EPHA4* | Chinese | De novo | c.2546G>A | p.Cys849Tyr | Missense | 1 | 28.2 | 0 | 0 |
| SCO2003P2080 | 2 | *EPHA4* | Chinese | De novo | c.1318+10344A>G | NA | Intronic | 1 | NA | 0 | 0 |
| SCO2003P3202 | 2 | *EPHA4* | Chinese | De novo | 2q35-36.2<br>4.6 Mb deletion | NA | CNV | NA | NA | 0 | 0 |
| SCO2003P3332 | 2 | *NGEF* | Chinese | De novo | c.1A>G | p.Met1? | Start lost | 0.95 | 12.8 | 0 | 0 |
| TSRHC01 | 2 | *NGEF* | Non-Hispanic<br>White | AD | c.857C>T | p.Ala286Val | Missense | 0.95 | 29.6 | 0.00002 | 0 |

Abbreviations: Chr (chromosomal localization); AA (amino acid); AD (autosomal dominant); CNV (copy number variant); ExAC PLI (probability of being loss-of-function intolerant from Exome Aggregation Consortium); CADD (Combined Annotation Dependent Depletion score); gnomAD (Genome Aggregation Database); NA (not applicable).

We next investigated *de novo* non-coding variants of *EPHA4* via whole genome sequencing (WGS) in 116 trio families with IS. A *de novo* heterozygous *EPHA4* intronic variant (c.1318+10344A>G) was identified in a female patient (SCO2003P2080) in an age range of 15-20 (Fig. 1D and Table 1). She developed scoliosis in her 10s with a 70° main curve Cobb angle. This variant is predicted to affect the branch point of the fifth intron of *EPHA4* (Fig. S2B). We performed nested PCR to show that this variant induced exon 5 skipping, resulting in a 339-bp in-frame deletion (Fig. 1H, J).

Next, we employed an in-house gene matching approach under the framework of DISCO study, which identified a patient (SCO2003P3202) in an age range of 5-10, who had been previously diagnosed with Waardenburg syndrome caused by a 4.46 Mb *de novo* deletion at 2q35-q36.2 (Li et al., 2015) (Table 1). Intriguingly, the patient also presented mild scoliosis (Fig. 1E). The deletion included the entire *PAX3* gene, which was responsible for the Waardenburg syndrome phenotype, and 36 neighboring genes, including *EPHA4* (Fig. 1K). As scoliosis is not typically associated with Waardenburg syndrome caused solely by *PAX3* pathogenic variants (Tassabehji et al., 1993), we hypothesize that the deletion of *EPHA4* may be responsible for the IS phenotype in this patient. Notably, the patient IDs used in this study are unique to our research group and will not be recognizable to anyone outside the research team, including hospital staff and the patients themselves.

Notably, the GWAS signal which we mapped to *EPHA4* (rs13398147) (Zhu et al., 2015) represents a significant eQTL in esophagus and colon tissues, with the T allele associated with decreased expression of *EPHA4*. In our East Asian GWAS cohort of 6,449 adolescent IS patients and 158,068 controls, we identified another two eQTLs in *EPHA4* associated with decreased expression of *EPHA4* in brain tissue (Table S3). In the same GWAS cohort, common SNPs in *EPHA4*, after aggregation, also showed significant enrichment (P=0.023) in IS patients versus controls. Taken together, the convergence between rare and common variants of *EPHA4* that lead to LoF or hypomorphic effects highlights the pivotal role of *EPHA4* in the pathogenesis of IS.

### IS-like phenotypes in zebrafish *epha4* mutants

To investigate the role of *EPHA4* in scoliosis, we utilized zebrafish as a model system due to its versatile nature in modeling adolescent IS (Bagnat and Gray, 2020; Boswell and Ciruna, 2017; Grimes et al., 2016; Xie et al., 2022). Zebrafish have two homologs of *EPHA4*, *epha4a* and *epha4b*. Using CRISPR-Cas9, we established a stable *epha4a* zebrafish mutant line with a 63-bp deletion in exon 3, which introduced a stop codon and resulted in a truncated protein (Fig. 2A-B). The homozygous *epha4a* mutant larvae had no apparent defects in either notochord or body axis development (Fig. S3A). Consistent with previous reports, the hindbrain rhombomeric boundaries were found to be defective in both the *epha4a* homozygous mutants and morphants (Fig. S3B) (Cayuso et al., 2019; Cooke et al., 2005). Interestingly, more than 75% adult mutants showed mild scoliosis (88 of 116), and some mutants exhibited severe scoliotic phenotype (4 of 116) (Fig. 2C-D, Movie S1&S2). Remarkably, some heterozygous adult mutants also developed mild scoliosis (14 of 95), whereas none of the wild-type fish showed any signs of scoliosis (0 of 76) (Fig. 2C-D). Similarly, we further generated the *epha4b* mutants with a 25-bp deletion in exon 3, which resulted in a frameshift mutation (Fig. S4A). The *epha4b* zebrafish mutants also developed mild scoliosis (28 of 43) (Fig. S4B-C). Intriguingly, both *epha4a* and *epha4b* mutants exhibited early onset scoliosis starting from around 20 days post fertilization (Fig. S4D), a stage similar to that of IS patients. Collectively, these data suggest that mutations in Epha4 proteins are linked to the scoliotic phenotype in zebrafish.

**Figure 2.**
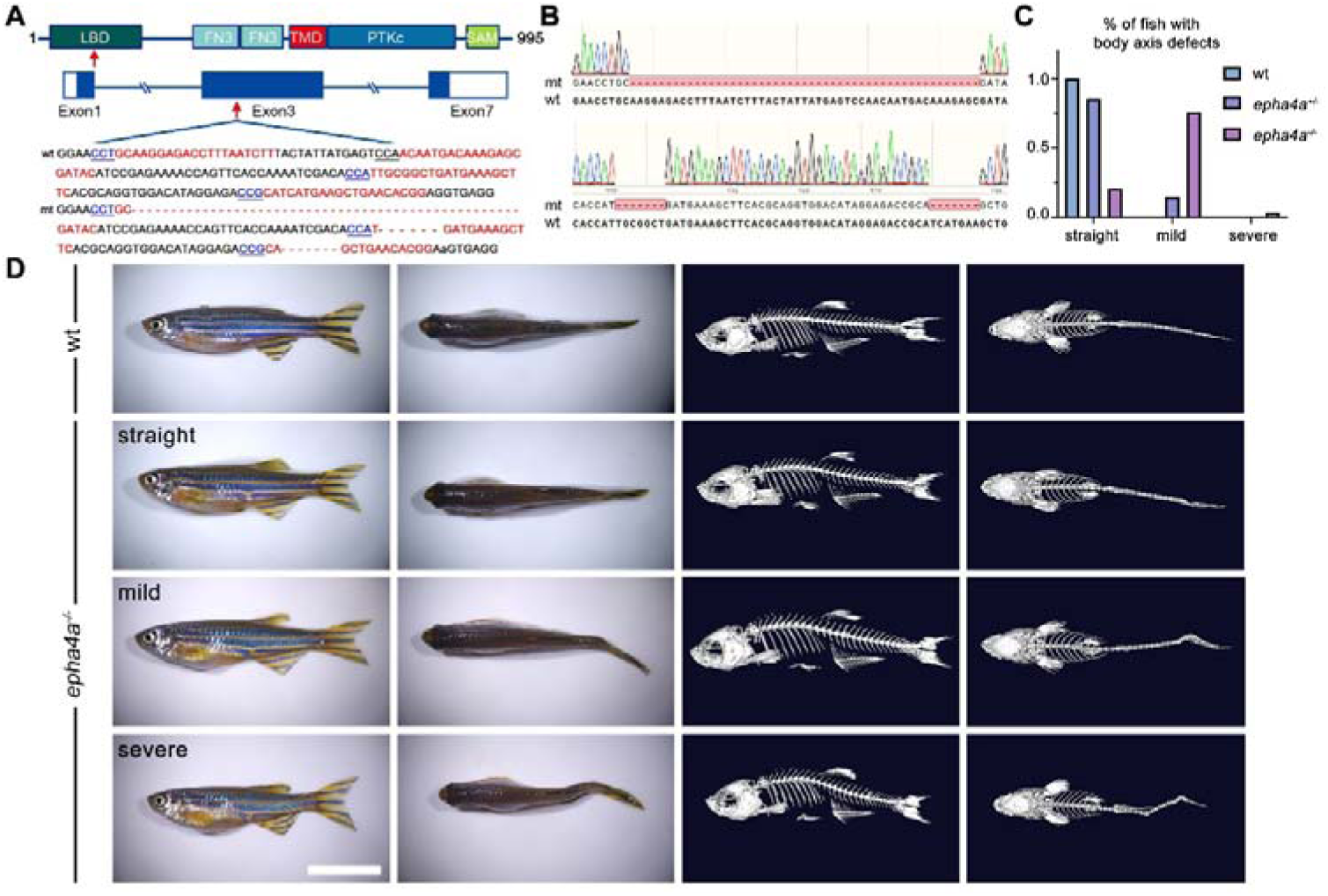
Scoliosis in zebrafish *epha4a* mutants. **A:** Diagram of the protein domains, genomic structures, and sequences of wild-type and corresponding *epha4a* mutants. Red arrows indicate mutation sites. Blue boxes indicate open reading frames. Underlined sequences indicate the protospacer adjacent motif (PAM) region, and red fonts indicate Cas9 binding sites. LBD: ligand binding domain; FN3: fibronectin type 3 domain; TMD: transmembrane domain; PTKc: catalytic domain of the protein tyrosine kinases; SAM: sterile alpha motif. **B:** Sanger sequencing results confirmed the deletion of the target region in *epha4a* mutant transcripts. **C:** Bar graph showing the percentages of adult zebrafish with normal, mild, or severe body axis defects in wild-type (n=76), *epha4a* heterozygote (n=95), or *epha4a* homozygous (n=116). **D:** Representative images of wild-type and homozygous *epha4a* mutants. Micro CT images are shown on the right. Lateral and dorsal views are shown. Scale bar: 1 cm.

### Abnormal left-right swimming pattern in the absence of Epha4a

While the *epha4a* mutants seemed to be grossly normal during the larvae stages, they developed spinal curvature gradually during later development. We decided to investigate whether these mutants exhibited any abnormalities in their behavior at the larvae stages using EthoVision XT software. By monitoring the swimming behavior of 8 days post-fertilization (dpf) zebrafish larvae, we found the motion distance and swimming velocity were significantly decreased in *epha4a* mutants (Fig. 3A-B). In addition, we observed a remarkable difference in the relative turning angle and angular velocity between these two groups. The wild-type group changed their swimming direction randomly, showing a relative angle around 0 degrees (Fig. 3C-D). In contrast, the *epha4a* mutants favored turning to one side of their directions (a positive angle reflects a leftward turn) (Fig. 3C-D). Moreover, the absolute turn angles and turning speed (angular velocity) were significantly higher in these mutants (Fig. 3E-F). To rule out potential off-target effects, we injected *epha4a* mRNA into *epha4a* mutants, which significantly restored the swimming coordination defects (Fig. S5).

**Figure 3.**
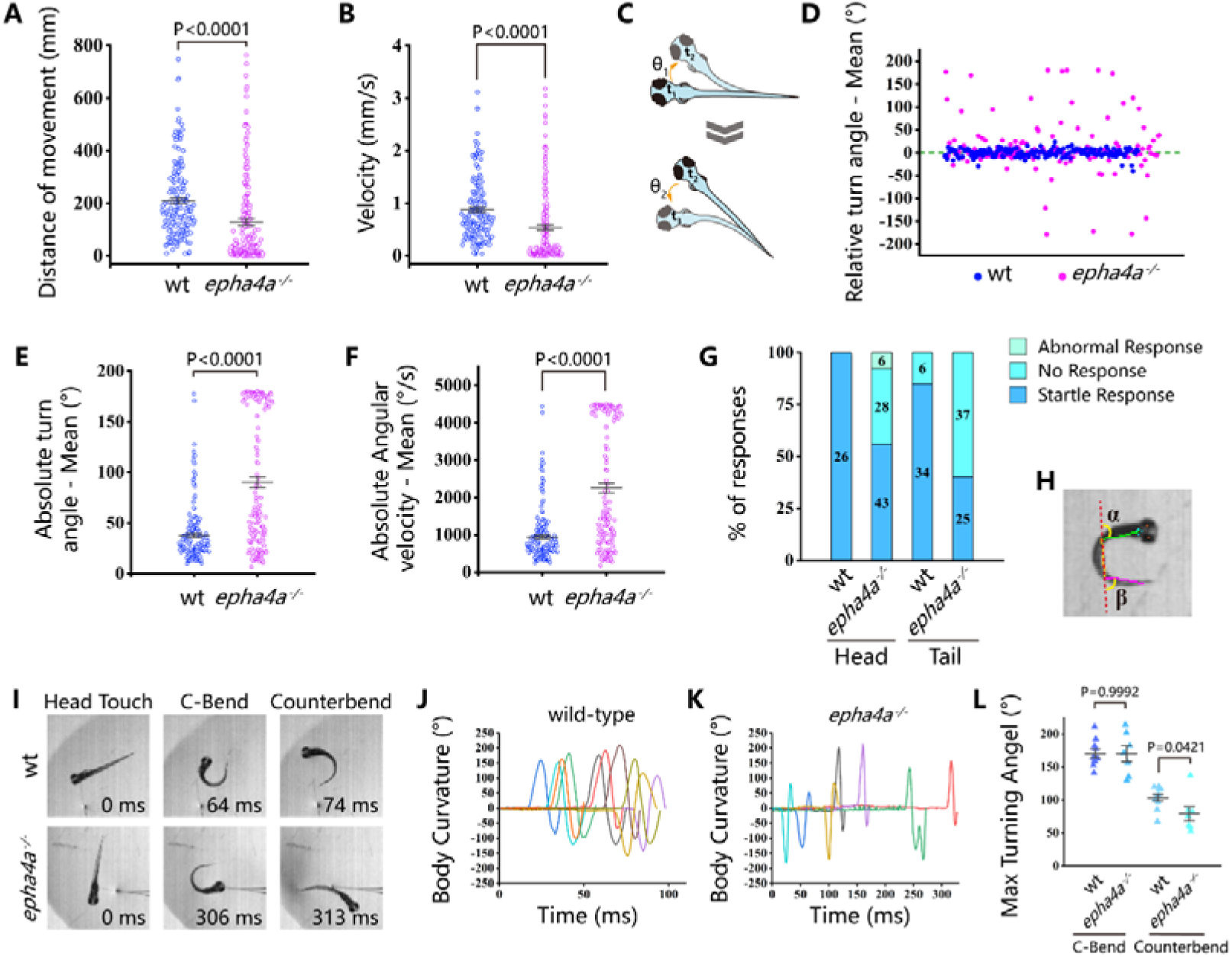
Abnormal left-right swimming pattern in the absence of Epha4a. **A:** Dot plots showing the swimming distance of each 8 dpf larva at a duration of 4 min (N=54 for wild-type and N=60 for *epha4a* mutants). **B:** Dot plots showing the swimming velocity of wild-type and mutant larvae as indicated. **C:** Diagram showing the turning angle (θ) of the larvae during swimming. **D:** Scatter plot showing the relative turning angle of wild-type and mutant larvae. The relative angles were calculated by the sum of turning angles during fish swimming with left (positive) or right (negative) turns. The *epha4a* mutants favored turning to one side of their directions compared with those of wild-type larvae. **E:** Dot plots showing the average absolute turning angle of wild-type and mutant larvae as indicated. **F:** Dot plots showing the average absolute angular velocity of wild-type and mutant larvae as indicated. **G:** Bar graph showing the percentages of 5 dpf zebrafish larvae with different reactions after tactile stimulation. N=10 for each group; the numbers of tactile stimulations are indicated in each column. **H:** Representative images of the total body curvature measurements in zebrafish larvae, with values as the sum of α and β angles shown in the figure. **I:** Representative time-series images of 5 dpf wild-type and *epha4a* mutant zebrafish larvae after tactile stimulation to the head. Each panel represents the points of maximal body curvature for the C-bend and counterbend after the tactile startle response. **J, K:** A plot of body curvature angles as measured in panel (H) during swimming in response to tactile head stimulation in 5 dpf wild-type (Panel J, N=5 larvae, n=10 stimuli) and *epha4a* mutants (Panel K, N=5 larvae, n=7 stimuli). Each colored curve represents an independent experiment showing the response of a single larva to a stimulus. The positive angle means turning right. **L:** The maximum curvature angles during the first C-bend and counterbend after tactile stimulation in wild-type and *epha4a* mutant larvae.

Next, we compared the swimming behavior after startle response between wild-type and *epha4a* mutants. We used a needle to touch the head or the tail of 5 days post-fertilization (dpf) zebrafish larvae to stimulate swimming behavior. In sibling controls, the larvae responded to the tactile stimulation and swam away quickly (Movie S3). Conversely, the *epha4a* mutants failed to respond to the initial stimulation, and the swimming pattern was defective with an abnormal bending pattern (37 of 62 from tail-stimulation group and 34 of 77 from head-stimulation group) (Fig. 3G, Movie S4). We further performed more comprehensive analysis by high-speed video microscopy. After tactile stimulation, wild-type larvae displayed a high-speed C-bend turn followed by a weaker counterbend turn after several milliseconds (Fig. 3H-I). This rhythmic left/right swimming pattern ensures that the fish swim away from the frightening stimulus. The turning angles of the control larvae after stimulation changed with a sinusoidal wave pattern (Fig. 3J). In contrast, this pattern was dramatically different in *epha4a* mutants (Fig. 3I,K). Of note, although the C-bend turning angles were similar between *epha4a* mutants and control siblings, the turning angles of the counterbend decreased significantly (Fig. 3L), implying a left/right coordination defect. Taken together, both regular swimming and tactile stimulation analyses suggested that the left-right coordination swimming pattern is compromised in the absence of Epha4a.

### Defects of left-right coordination due to abnormal CPG in the absence of Epha4a

The coordinated left-right locomotion of zebrafish larvae relies on the synchronized contraction of muscle fibers, a process regulated by motor neurons situated on each side of the fish. To explore this intricate mechanism, we utilized a Tg(*elavl3*:GAL4; *UAS*: GCaMP) double transgene, allowing the expression of a genetically encoded calcium sensor in all neurons. We observed the rhythmic activation of calcium signaling in motor neurons located within the spinal cord (Fig. 4A, Fig. S6A, Movie S5, Movie S6). In wild-type larvae, the calcium signals exhibited an alternating pattern between the left and right sides of the body, whereas this coordinated pattern was disrupted in both *epha4a* mutants and morphants (Fig. 4A-B vs A’-B’, Fig. S6A-B vs A’-B’, Movie S7, Movie S8). Additionally, we found that the activation frequency of motor neurons on the left and right sides was comparable in wild-type larvae, but significantly different in the absence of Epha4a (Fig. 4C-D, Fig. S6C-D).

**Figure 4.**
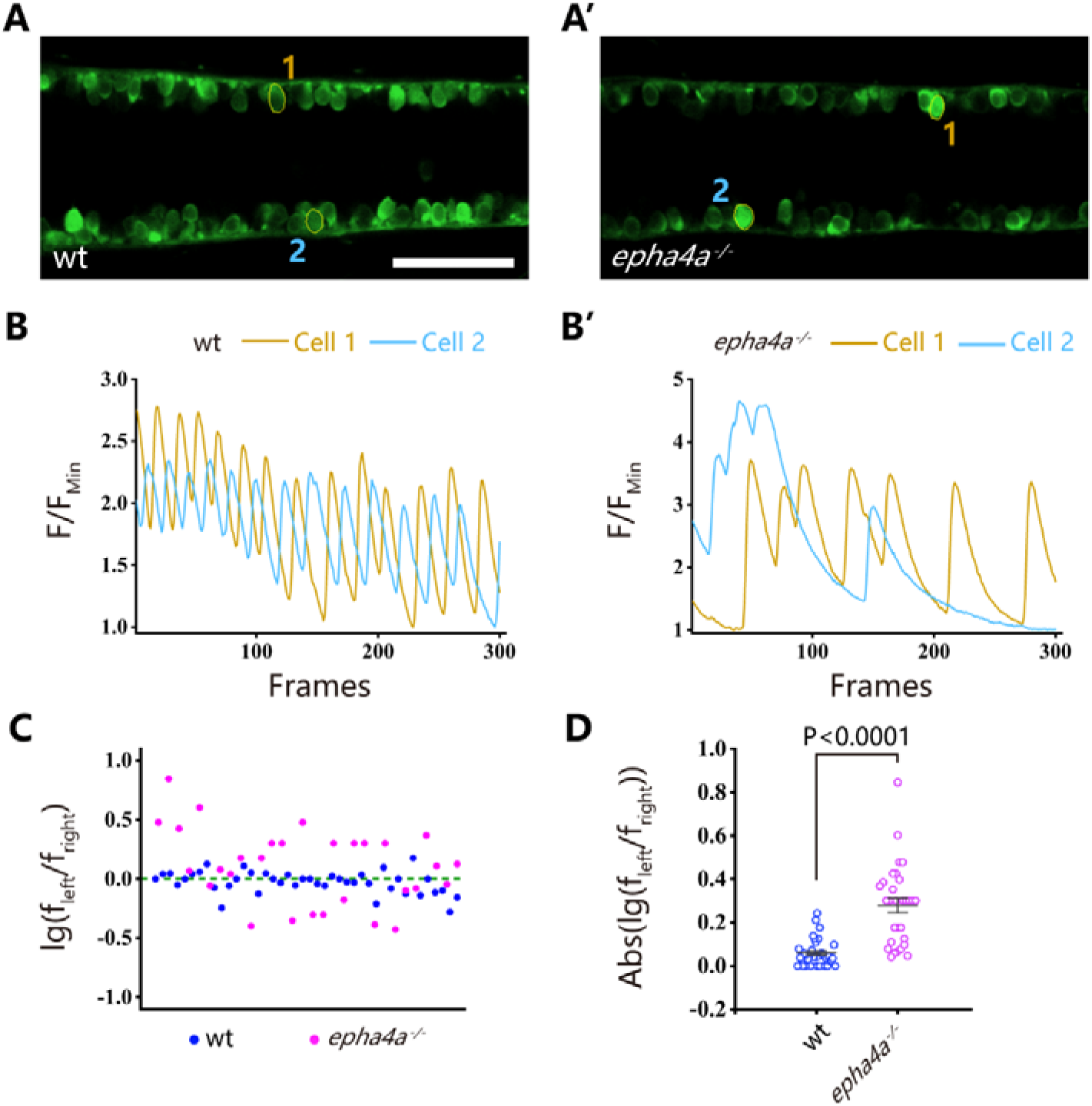
Uncoordinated left-right activation of spinal cord neurons in *epha4a* mutants. **A, A’:** Fluorescent images showing the dorsal-view of 24 hpf Tg(*elavl3*:GAL4; *UAS*:GCaMP) double transgenic larvae. The corresponding movies are shown in Movie S5 and S6. **B, B’:** Line charts showing the quantification of fluorescence changes of the region of interests (ROIs, circled in A, A’) in wild-type larvae and *epha4a* mutants. **C:** Scatter plot showing the distribution trend of the ratio of the calcium signal frequency between left and right in wild-type (N=14 larvae, n=42 experiments) and *epha4a* mutants (N=10 larvae, n=30 experiments). **D:** Statistical graph of the ratio of the calcium signal frequency between left and right in wild-type larvae and *epha4a* mutants. Scale bars: 50 μm in panel **(A, A’)**.

One well-established concept regarding left-right coordination involves the presence of central pattern generators (CPGs), which are regulated by interneuron circuitry within the spinal cord. We further examined the axon guidance of interneurons in *epha4a* mutants. First, we investigated the commissural trajectories of reticulospinal (RS) interneurons. In control larvae, the large Mauthner neurons, along with other RS neurons, were symmetrically positioned on each side of the midline and projected their axons to the contralateral sides (Fig. 5A). These bilaterally projected axons typically crossed at the midline and subsequently synapsed on motoneurons of the opposite sides, contributing to the generation of spinal cord neural circuits (Hale et al., 2016). However, in *epha4a* mutant larvae, we observed an abnormal pattern in axonal projections. Specifically, the mutant axons failed to traverse the midline and instead extended ipsilaterally (Fig. 5A). In addition, the distance between Mauthner neurons (rhombomere 4) and rhombomere 7 was significantly decreased and rhombomere 5 was scarcely visible in the mutant larvae (Fig. 5A-B). Furthermore, the sites of axon crossing between two Mauthner neurons tended to deviate to one side of the midline in the mutants (Fig. 5C-D’).

**Figure 5.**
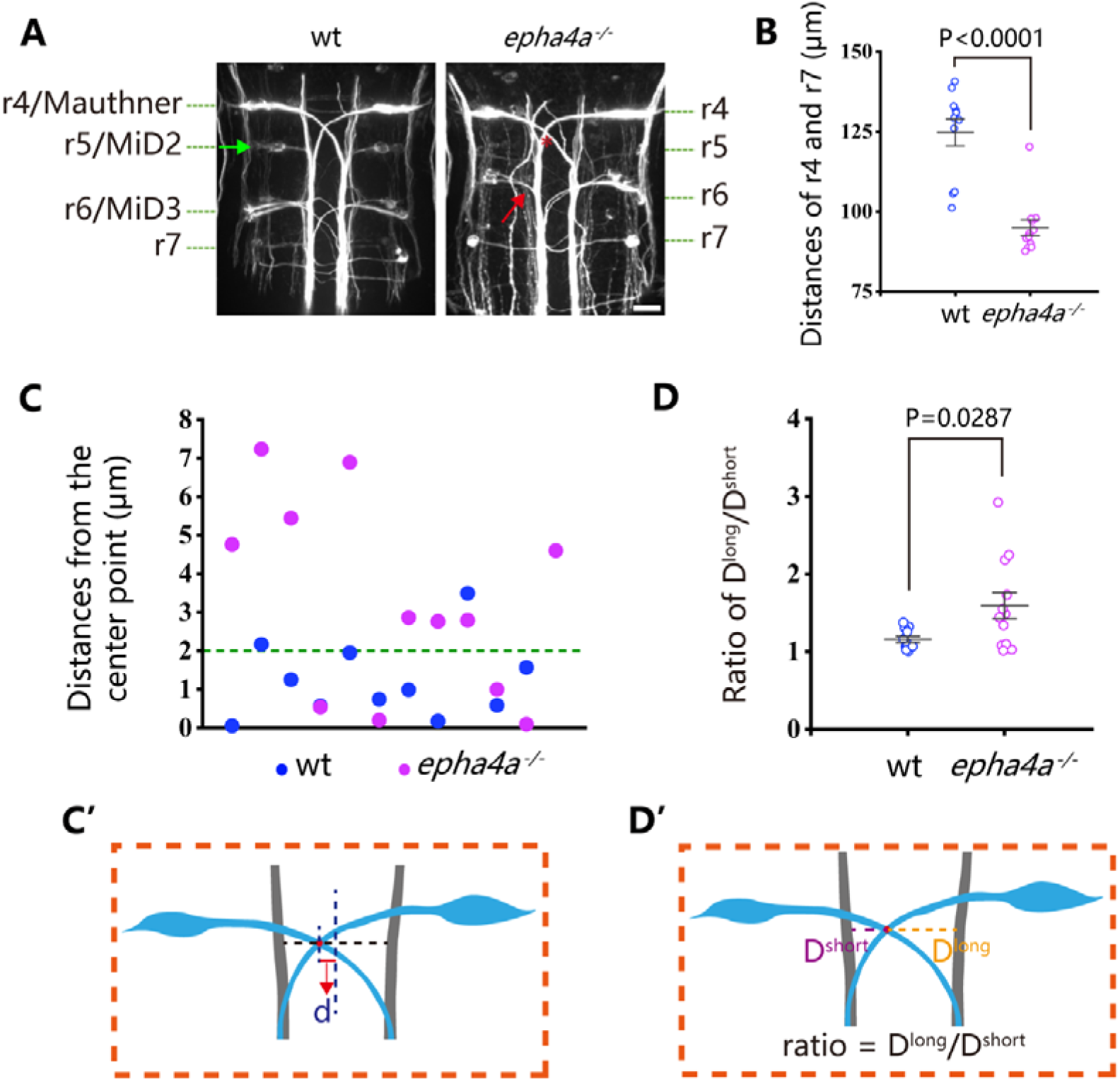
Disorganized neural patterning in *epha4a* mutants. **A:** Confocal images showing reticulospinal neuronal axons in 48 hours post-fertilization (hpf) wild-type and *epha4a* mutant larvae visualized with anti-neurofilament antibody RMO44. Asterisks indicate the cross sites of Mauthner axons. The green arrow indicates the cell body of the r5/MiD2 neuron in a wild-type larva. The red arrow points to the ipsilaterally projected axon of r6/MiD3 in the mutant larva, which is normally projected to the other side in wild-type fish. **B:** Statistical chart showing the distance between r4 and r7 of 48 hpf wild-type and *epha4a* mutants. **C:** Scatter plot showing the distance (d) between the center line and the intersection site of Mauthner axons as indicated in panel C′. **D:** The ratio of the distance between the intersection site of Mauthner axons and bilateral axon bundles in 48 hpf wild-type (N=11 larvae) and *epha4a* mutants (N=12 larvae). The ratios were calculated as in panel D′. Scale bars: 20 µm in panel **(A)**.

Cerebrospinal fluid-contacting neurons (CSF-cNs) represent a unique type of interneuron responsible for modulating the V0 and V2a interneurons, which are integral components of locomotor CPGs (Fidelin et al., 2015; Talpalar et al., 2013; Wu et al., 2021). In wild-type larvae, we observed that the ascending axons of these neurons projected either to the right or left side from the midline, as visualized using Tg(*pkd2l1*:GAL4;*UAS*:Kaede) double transgenic larvae (Fig. 6A). However, in the absence of Epha4a, the projection of these neurons exhibited notable disorganization, with numerous axons crossing the midline from one side of the trunk. This disorganized pattern was observed in both heterozygotic and homozygotic mutants (Fig. 6A-B). Additionally, we employed an optogenetic approach to activate these CSF-cNs, utilizing the Tg(Gal4^s1020t^; *UAS*:ChR2) double transgene. Following optical stimulation, we observed robust tail oscillations as previously described (Wyart et al., 2009). In wild-type larvae, these tail oscillations exhibited a symmetrical left-right beating pattern(Fig. 6C-D, Movie S9). However, a striking disruption of this symmetry was observed in *epha4a* morphants, as they consistently beat towards one side of the trunk following optical stimulation (a positive angle reflects a rightward turn) (Fig. 6C’-D’ and E, Movie S10). Collectively, these findings demonstrate that the left-right coordination deficiencies observed in *epha4a* mutants arise from abnormal neural circuit formation, which consequently disrupts the integrity of the CPGs.

**Figure 6.**
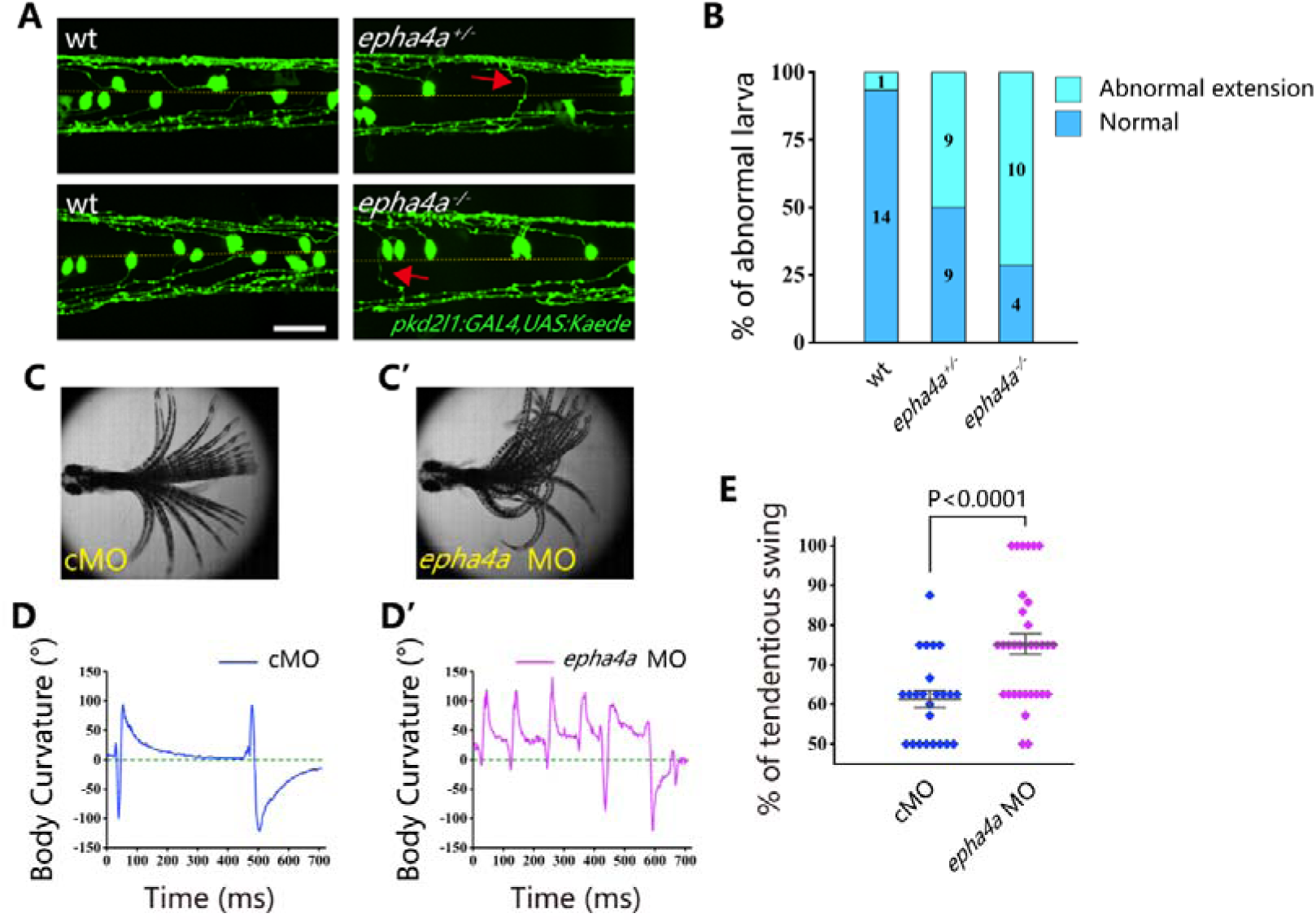
Aberrant swimming as a result of abnormal extension of CSF-cNs axons. **A:** Fluorescent images showing the distribution of ascending axons of CSF-cNs marked by Tg(pkd2l1:GAL4,UAS:Kaede) in 2 dpf *epha4a* mutant larvae. Yellow line indicates the midline and the red arrows indicate aberrantly extended axons in *epha4a*^+/-^ and *epha4a*^-/-^ larvae. **B:** Bar graph showing the percentages of abnormal extension of CSF-cNs axons in 2 dpf wild-type, *epha4a* heterozygote, and *epha4a* homozygous. The numbers of larvae are indicated in each column. **C, C’:** Superimposed frames of tail oscillations in 5 dpf control and *epha4a* morphants. **D, D’:** A plot of body curvature angles in panel (C) and (C’). The positive angle means turning right. **E:** Percentages of tendentious swing in control (N=8 larvae, n=24 experiments) and *epha4a* morphants (N=11 larvae, n=33 experiments). The percentages were calculated by the ratio of tendentious tail oscillation during the first eight swings. Scale bars: 50 µm in panel **(A)**.

### Ephrin B3**-**Epha4 signaling regulates interneuron axon extension

To further explore the role of Epha4a during interneuron axon extension, we examined the expression of *epha4a* during early zebrafish embryonic development. Whole-mount in situ hybridization results showed that both *epha4a* and *epha4b* were abundantly expressed in the zebrafish spinal cord (Fig. S7A). We plotted the expression of these two genes using two published single-cell transcriptome data, which showed that *epha4a* and *epha4b* were both expressed in interneurons, suggesting a role for *epha4* in interneuron function (Fig. S7B) (Cavone et al., 2021; Scott et al., 2021). Notably, the expression of *efnb3b*, encoding the ligand of Epha4, was highly enriched in the midline floor plate cells (Fig. S7B).

Ephrins, through interacting with Eph receptors, play a critical role in repulsive axon guidance during neural development (Egea and Klein, 2007; Flanagan and Vanderhaeghen, 1998). We further analyzed axon guidance in *efnb3b* morphants. Similar to those of *epha4a* mutants, the *efnb3b* morphants also displayed axon guidance defects, as well as uncoordinated calcium activation (Fig. S8A-C). In addition, morphants larvae also displayed left-right oscillations defects after optogenetic stimulation (a positive angle reflects a rightward turn) (Fig. S8D-E).

### Candidate variants in the *EPHA4*-related genes

Our zebrafish studies suggested that EPHA4 signaling is crucial for interneuron axon guidance, hence the formation of functional CPGs. Next, we further asked whether mutation of other components of the EPHA4 signaling can result in IS in humans. By searching for rare variants in *EPHA4*-related genes (Fig. S9) (Szklarczyk et al., 2019), we identified heterozygous *de novo* start-loss variant (c.1A>G, p.Met1?) in *NGEF* in a young male (SCO2003P3332) (Fig. 7A-B, Table 1), whose scoliosis was diagnosed in an age range of 15-20, with a main curve Cobb angle of 60°. *NGEF* encodes the neuronal guanine nucleotide exchange factor Ephexin that differentially affects the activity of GTPases RHOA, RAC1, and CDC42. The activation of Ephexin is triggered by ephrin through EPHA4 (Shamah et al., 2001).

**Figure 7.**
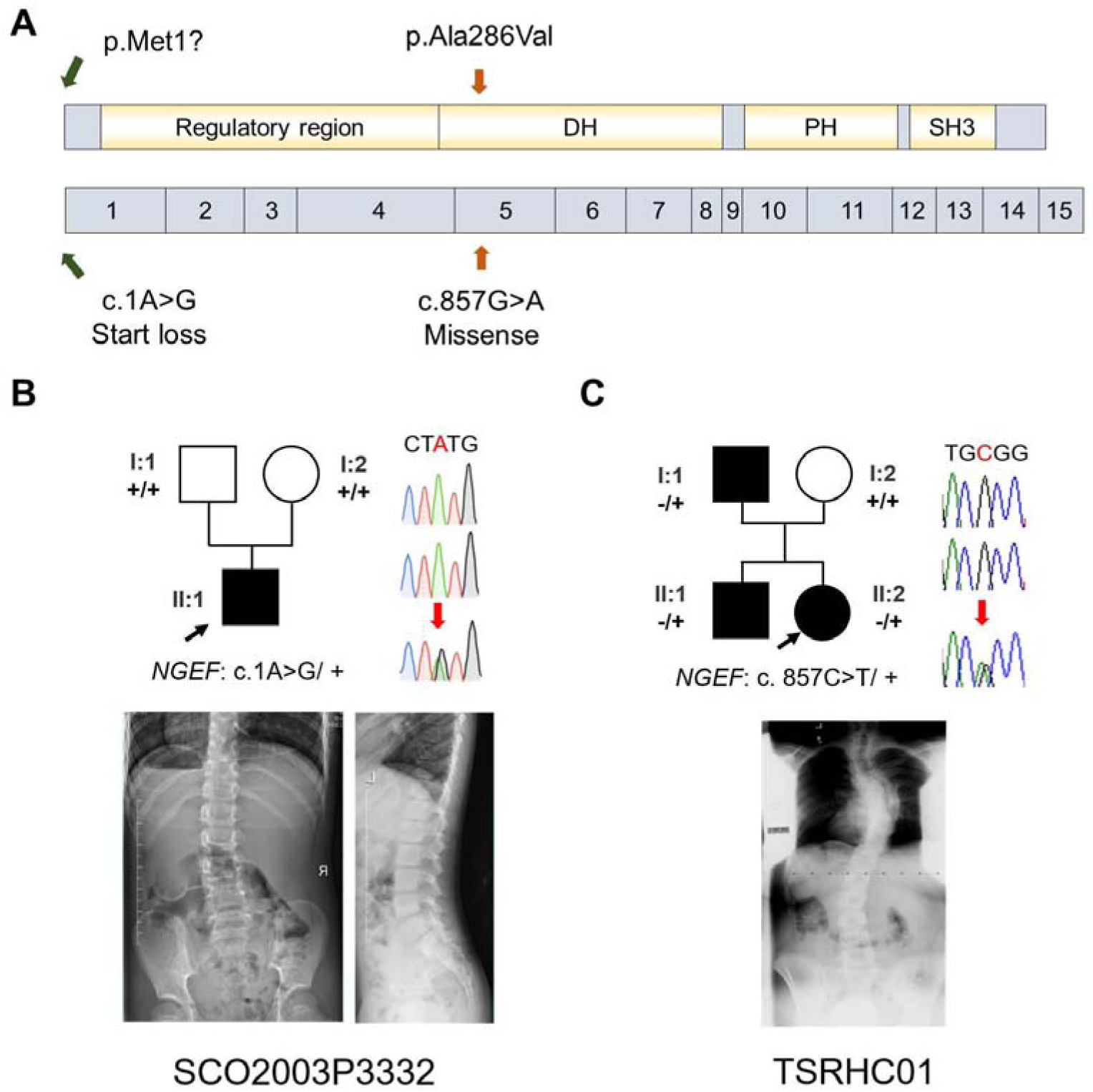
IS patients with potential *NGEF* variants. **A:** Protein structure of NGEF protein with the position of potential variants. **B, C:** Pedigrees and spinal radiographs of two probands with dominant gene variants. Sanger sequencing results are shown on the right.

Strikingly, we further identified a dominant missense variant (c.857G>A, p.Ala286Val) in *NGEF* in a quad family with three affected members (Fig. 7C, Table 1). The proband (II:2, TSRHC01) has a 58° main curve Cobb angle. This variant, which is located in the RhoGEF domain of Ephexin, is predicted to be highly deleterious (CADD=29.6).

Altogether, our results suggest that defects of the CPGs owing to abnormal EPHA4 signaling maybe one of the crucial factors responsible for IS.

## Discussion

IS is a disease with diverse causes, and the underlying mechanisms can vary even among patients with similar scoliotic phenotypes. Previous genetic studies have highlighted the significance of the extracellular matrix (ECM) in maintaining the balance of axial bone and supporting soft tissues in the spine, thus playing a crucial role in IS development (Haller et al., 2016). For example, the top SNP associated with IS maps to *LBX1*, an essential molecule for ECM maintenance and bone homeostasis (Takahashi et al., 2011). Additionally, genetic loci in muscle development-related genes have also been associated with the onset of scoliosis, emphasizing the intricate interaction between bones and muscles (Ogura et al., 2015). However, it is worth noting that the variants in these genes explain only a small fraction of the overall heritability of IS. Consequently, it is imperative to establish connections between the extensive genetic findings and biological mechanisms that can elucidate the etiological landscape of IS.

In this study, we mapped 41 significant genome-wide loci to functional genes through positional mapping and functional mapping such as eQTL. Then we determined the enrichment in patient cohorts of rare variants in these genes, which may have greater impacts compared with common SNPs. This approach revealed the convergence of SNPs and rare variants in *EPHA4* that are enriched in patients with IS. We also identified additional high-impact variants in *NGEF*, which is involved in the EPHA4 pathway.

Our studies using zebrafish have revealed that deficiency of Epha4 can lead to the development of scoliosis. Interestingly, we observed that even heterozygotic *epha4a* zebrafish mutants displayed mild scoliosis (Fig. 2C), which is consistent with the occurrence of this condition in scoliosis patients. Further analysis involving behavior and imaging demonstrated that the absence of Epha4a resulted in defective left-right coordination. This coordination is crucially governed by CPGs, which generate rhythmic patterns of neural activity to coordinate limb movements on both sides of the body (Kiehn, 2006; Marder and Bucher, 2001; Talpalar et al., 2013). Previous studies have reported the involvement of ephrin and its receptors in axon guidance during the maturation of neural circuits, including CPGs (Andersson et al., 2012; Borgius et al., 2014; Iwasato et al., 2007). Ephrin B3, present in radial glial cells at the dorsal midline, serves as a repulsive barrier to axons expressing Epha4 (Imondi et al., 2000). Axon guidance is disrupted when this repulsive pathway is inactivated, leading to midline invasion (Butt et al., 2005; Kullander et al., 2001; Paixão et al., 2013). In *Epha4* and *Efnb3* knockout mice, a ‘hopping’ gait is observed, which is attributed to the aberrant crossing of motor neurons expressing Epha4 (Kullander et al., 2003). Consistent with these observations, we observed disrupted axon guidance in the interneurons, which are integral components of CPGs, in both *epha4a* mutants and *efnb3b* morphants. This finding suggests that CPG malfunction may be a significant contributing factor to the scoliosis phenotype observed in these genetic backgrounds. It is highly likely that the lack of coordinated left-right locomotion generates imbalanced mechanical forces on the spine, gradually leading to spinal curvature during later stages of development (Fig. 8).

**Figure 8.**
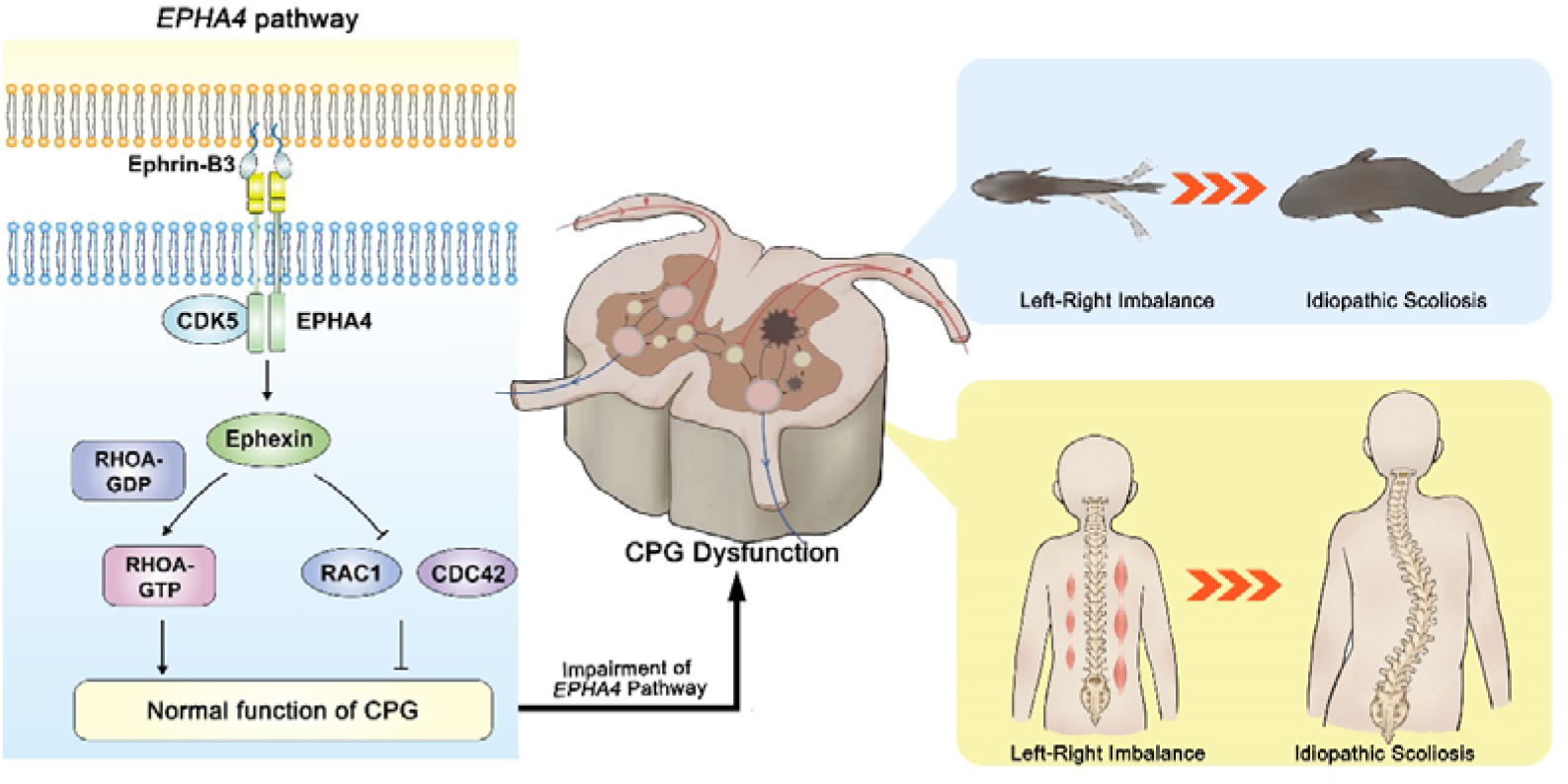
The proposed mechanism of IS mediated by *EPHA4* dysfunction. In healthy individuals, EphrinB3-activated EPHA4 phosphorylates CDK5, leading to the phosphorylation of Ephexin, a protein encoded by NGEF. Phosphorylated Ephexin can regulate axon guidance through either activating RHOA or suppressing CDC42 and RAC1 signaling. These processes are critical to maintain the normal function of the CPG, the local neural network that provides coordinated bilateral muscle control. Impairment of the EPHA4 pathway and CPG may cause an imbalance of the motor drive from the spinal cord during development, thus causing the uncoordinated left/right swimming behavior in zebrafish larvae and the asymmetry of the bilateral muscular pull in a young child. Although the appearance is normal in early childhood, the dysfunction produces a scoliotic curve during the growth spurt.

Thus, our data provided a novel biological mechanism of IS, i.e., the impairment of neural patterning and CPG. Previous studies have provided clues on the role of CPGs in IS. Patients with adolescent IS showed asymmetric trunk movement during gait, as characterized by increased relative forward rotation of the right upper body in relation to the pelvis (Kramers-de Quervain et al., 2004; Nishida et al., 2017). An electromyography (EMG) study also showed asymmetric activation of paraspinal muscles between the convex and concave sides at the scoliosis curve apex (Shimode et al., 2003). In a child with a strong family history of IS, asymmetric hyperactivity was observed by EMG months before scoliosis was evident (Valentino et al., 1985). These left-right locomotor coordination abnormalities indicated the maldevelopment of CPGs as potential cause of IS. The CPG asymmetry may induce an imbalance in trunk muscle strength, resulting in asymmetric rib drooping. This leads to an abnormal vertebral rotation and then the onset of IS. Moreover, in a companion study, Wang et al. identified a number of rare variants in *SLC6A9*, which encodes glycine transporter 1 (GLYT1), in familial and sporadic adolescent IS cases. The *slc6a9* mutant zebrafish also exhibited discoordination of spinal neural activities with pronounced lateral spinal curvature, recapitulating the human IS phenotype (Wang et al., 2024). Taken together, we propose that the dysfunction of CPGs would cause an imbalance in the motor drive from the spinal cord and the asymmetric transversospinalis muscle pull, eventually producing a scoliotic curve (Fig. 8).

In summary, our study demonstrates that both common and rare variants within the EPHA4 pathway contribute to the genetic architecture of IS. EPHA4 pathway dysfunction causes axon pathfinding defects, resulting in impaired coordinated left-right locomotion by disrupting neural patterning and the function of CPGs, thereby potentially leading to IS.

## Materials and Methods

### Mapping of candidate genes utilizing previous association studies

Through a systematic literature review, we identified IS-associated SNPs reported in GWASs and meta-analyses of GWAS. The literature search was carried out using MEDLINE (via Pubmed.gov) and Web of Science (via Clarivate Analytics) and was limited to English-language articles published from January 1980 to October 2020. The following keywords were combined to perform the search: ‘idiopathic scoliosis’ AND ‘GWAS’ OR ‘SNP’ OR ‘single nucleotide polymorphism’ OR ‘variant’ (Table S4). The inclusion/exclusion criteria of abstracts are provided in Table S5. After screening the titles and abstracts, we obtained from the full-text articles the rsID, chromosome, and position for SNPs with a threshold for genome-wide significance of P < 5.0 × 10^−8^. For SNPs identified in multiple studies, we recorded the lowest P-value.

SNPs reported in previous studies were first pruned using Functional Mapping and Annotation (FUMA, v1.4.1, https://fuma.ctglab.nl/) (Watanabe et al., 2017). Significant SNPs were considered independent at r^2^ < 0.6. All known SNPs (available in the 1000 Genomes reference panel, https://www.internationalgenome.org/) were included for further gene mapping if they were in a linkage disequilibrium (LD) block (r^2^ ≥ 0.6) with a significant independent SNP. Three SNP-to-gene strategies were used:

1. For positional mapping of significant independent SNPs, we used annotations obtained from ANNOVAR (http://annovar.openbioinformatics.org). A 10-kb maximum distance was applied for intergenic SNPs.
2. For the eQTL mapping, we mapped significant independent SNPs and SNPs in an LD block to eQTLs acorss 44 GTEx tissue types (release V8) (Consortium, 2020). SNP-gene pairs with a false discovery rate (FDR) ≤ 0.05 were considered significant.
3. For chromatin interaction mapping, we overlapped the significant independent SNPs and those in LD blocks with one end of significantly interacting regions across various tissue and cell types. Information on the interacting regions was derived from Hi-C datasets of 21 tissues and cell types provided by GSE87112. The significance of interactions for Hi-C datasets was computed by Fit-Hi-C, with an FDR ≤ 10^-6 considered significant. Genes were mapped if their promoter regions overlapped with another end of the significant interactions. The promoter region was defined as the region from −250 bp to +500 bp relative to the transcription start site.

### Cohort description

The Peking Union Medical College Hospital (PUMCH) cohort: The PUMCH cohort comprised 411 unrelated Chinese patients with severe IS (Cobb angle ≥ 40°) who underwent spinal surgery in the PUMCH between October 2017 and March 2022 as part of the Deciphering disorders Involving Scoliosis and COmorbidities (DISCO) study (http://www.discostudy.org/). The clinical diagnosis was confirmed using standing full-spine radiographs, three-dimensional computed tomography, and magnetic resonance imaging. These patients did not show any congenital or neuromuscular defect at the time of recruitment. The control cohort consisted of 3,800 individuals without observable scoliosis and with exome sequencing or genome sequencing performed at PUMCH for clinical or research purposes. Individuals with vertebrae malformation or congenital developmental defects were excluded. A patient with a 2q35-36.2 deletion, including the *EPHA4* gene, was also recruited through in-house gene matching of the DISCO study (Li et al., 2015). This patient was subsequently evaluated for scoliosis by examination and radiography.

The East Asian cohort: The East Asian cohort includes totally 6,449 IS patients and 158,068 controls from four independent datasets (Japanese dataset 1 (Kou et al., 2013; Takahashi et al., 2011): 1,261 cases, 15,019 controls; Japanese dataset 2 (Ogura et al., 2015): 878 cases, 21,334 controls; Japanese dataset 3 (Kou et al., 2019): 3,333 cases, 119,630 controls; Hong Kong dataset: 977 cases, 2,085 controls). The inclusion criteria for IS subjects was as same as our previous studies (Fan et al., 2012; Kou et al., 2019).

The Texas Scottish Rite Hospital for Children (TSRHC) cohort: We used a replication cohort of European-ancestry patients with IS from the TSRHC. Cases considered for inclusion in the study met criteria for a positive diagnosis of IS: lateral deviation from the midline greater than 15 degrees as measured by the Cobb angle method from standing spinal radiographs, axial rotation toward the side of the deviation and exclusion of relevant co-existing diagnoses.

### Blood sample collection

In the PUMCH cohort, genomic DNA samples were extracted from peripheral blood leukocytes of each subject using a QIAamp DNA Blood Mini Kit (Qiagen, Hilden, Germany), according to the manufacturer’s protocol. Purified DNA was qualified by Nanodrop 2000 (Thermo Fisher Scientific, Waltham, MA, USA) and quantified by Qubit 3.0 using the dsDNA HS Assay Kit (Life Technologies, Carlsbad, CA, USA). DNA samples were stored at 4°C until used.

In the East Asian and TSRHC cohort, genomic DNA was extracted from peripheral blood or saliva using a standard protocol.

### DNA sequencing and variant calling

In the PUMCH cohort, whole exome sequencing (WES) or whole genome sequencing (WGS) was performed on peripheral blood DNA from all individuals and available family members (Table S6). A SureSelect Human All Exon V6+UTR r2 core design (91 Mb, Agilent) was used for exon capture. The exomes were then sequenced on an Illumina HiSeq 4000 (Illumina, San Diego, CA, USA) according to the manufacturer’s instructions. For WGS, sequencing libraries were prepared using the KAPA Hyper Prep kit (KAPA Biosystems, Kusatsu, Japan) with an optimized manufacturer’s protocol. We performed multiplex sequencing using an Illumina HiSeq X-Ten sequencer (Illumina, San Diego, CA, USA). Variant calling and annotation were done in-house using the Peking Union Medical College Hospital Pipeline (PUMP) described previously (Chen et al., 2021; Zhao et al., 2021).

In the TSRHC cohort, subjects in 162 adolescent IS families were sequenced as part of the Gabriella Miller Kids First Pediatric Research Consortium (GMKF) at The HudsonAlpha Institute for Biotechnology (Huntsville, AL). In summary, DNA was normalized, sheared and then ligated to Illumina paired-end adaptors. The purified ligated DNA was amplified and exome sequencing was performed on the Illumina HiSeq X platform. The sample’s sequences were aligned into GRCh38 and genotypes were joint-called per each family by the GMKF’s Data Resource Center (DRC) at Children’s Hospital of Philadelphia following GATK best practices as detailed here (Mukhopadhyay et al., 2020). GMKF DRC’s alignment and joint genotyping pipelines are open source and made available to the public via GitHub (https://github.com/kids-first/kf-alignment-workflow and https://github.com/kids-first/kf-jointgenotyping-workflow).

### Rare variant association analysis of the candidate genes

To determine the contribution of rare variants in GWAS candidate genes to IS, we analyzed the gene-based mutational burden for 156 candidate genes. Only ultra-rare variants with a gnomAD population-max allele frequency ≤ 0.01% and a cohort allele count ≤ 3 were analyzed. Each variant was assigned a weight, and the mutational burden of a given gene was defined as the maximum weight value among all ultra-rare variants carried by the individual. The SAIGE-GENE+ package was used to determine the weighted mutational burden test for each gene (Zhou et al., 2022).

Weighting criteria for the weighted burden analysis were developed according to the variant types and in silico results. The LoF variants (canonical splicing variants, nonsense variants, or variants that cause frameshift, stop-gain, or start-loss) were calculated together with the protein-altering variants, including missense variants and in-frame indels. Each variant was assigned a weight range from 0-1 and the mutational burden for a given gene was defined as the maximum weight value among all ultra-rare variants carried by the individual. The LoF variants annotated as ‘high confidence’ by the loss-of-function transcript effect estimator (LOFTEE) (Karczewski et al., 2020) were assigned a weight value of 1.0. The LoF variants annotated as ‘low confidence’ or unlabeled by LOFTEE, the non-canonical splicing variants with a SpliceAI score > 0.5, and the missense variants with a rare exome variant ensemble learner (REVEL) (Ioannidis et al., 2016) score of 0.8 were assigned a weight value of 0.8. The in-frame insertions/deletions (indels) with a Combined Annotation Dependent Depletion (CADD) (Kircher et al., 2014) score > 20 and the missense variants with a REVEL score > 0.6 and ≤ 0.8 were assigned a weight value of 0.6. The in-frame indels with a CADD score > 10 and ≤ 20 and the missense variants with a REVEL score > 0.4 and ≤ 0.6 were assigned a weight value of 0.4. The in-frame indels and the missense variants with a REVEL score > 0.2 and ≤ 0.4 were assigned a weight value of 0.2. The remaining missense variants were assigned a weight value of 0.

### Sanger sequencing of familial participants

Sanger sequencing of familial participants was performed to determine the origin of variants in *EPHA4* and *NGEF*. All LoF variants and protein-altering variants identified in familial participants were validated. Variant-encoding gene regions were amplified by PCR from genomic DNA obtained from probands, as well as from parents for trios, to determine the origin of the variants. The amplicons were purified using an Axygen AP-GX-50 kit (Corning, NY, USA) and sequenced by Sanger sequencing on an ABI 3730xl instrument (Thermo Fisher Scientific, Waltham, MA, USA).

### Minigene assay

The splicing variant (*EPHA4:* c.1443+1G>C) was characterized by a minigene assay. Genomic DNA from the heterozygous patient was amplified by PCR using a high-fidelity DNA polymerase. Amplicons included exon 6, intron 6, exon 7, intron 7, and exon 8 of the *EPHA4* gene. PCR products were cloned into the vector via the restriction sites BamHI and MluI for pCAS2, which is based on the pcDNA3.1 plasmid (Thermo Fisher Scientific, Waltham, MA, USA). Clones with wild-type or mutant genomic inserts were selected and verified by sequencing the cloned DNA fragments. The recombinant plasmids were transfected into 293T cells using LipofectamineTM 3000 reagent (Thermo Fisher Scientific, Waltham, MA, USA). For RT-qPCR, total RNA was isolated from the transfected cells using Trizol reagent (Thermo Fisher Scientific, Waltham, MA, USA and reverse transcription was performed using the GoScript™ Reverse Transcription System (Promega, Madison, MI, USA). PCR amplification was performed using the pCAS2-RT-F and pCAS2-RT-R primers, and the products were sequenced using pCAS2-RT-F.

Primers sequences for the minigene assay were as follows:

pCAS2-RT-F: 5′-CTGACCCTGCTGACCCTCCT-3′

pCAS2-RT-R: 5′-TTGCTGAGAAGGCGTGGTAGAG-3′

### Nested PCR

The splicing variant (*EPHA4:* c.1318+10344A>G) was characterized by nested PCR. RNA was extracted from the whole blood of the patient using the TRIzon Reagent (CWBIO, Hangzhou, China) according to the manufacturer’s guidelines. The cDNA was synthesized using HiScript II 1st Strand cDNA Synthesis Kit with a gDNA wiper (Vazyme, Nanjing, China) according to the manufacturer’s guidelines. Nested PCR was performed as described previously (Yao and Tavis, 2005). Nested PCR reactions in 50 μl, included 2 μl cDNA from the reverse transcription reaction as the template for the first round of PCR or 2 μl of first-round PCR product as the template for the second PCR, 1.5 μl 10 μM sense primer, 1.5 μl 10 μM anti-sense primer, 5 μl nucleotide mix (2 mM each dNTP), 5 μl 10× KOD Buffer, 1 μl 1 unit/μl Kod-Plus-Neo polymerase, 3 μl 25 mM MgSO_4_ and 31 μl ddH_2_O. The PCR program was (94°C for 2 min, 98°C for 10 s, 55°C for 30 s, 68°C for 60 s) × 20 cycles and then (94°C for 2 min, 98°C for 10 s, 57°C for 30 s, 68°C for 60 s) × 30 cycles. Primers sequences:

For the first round:

F: 5′-GGCTCCTGTGTCAACAACTC-3′

R: 5′-GTTGGGATCTTCGTACGTAA-3′

For the second round:

F: 5′-AACTGCCTATGCAACGCTGG-3′

R: 5′-AGCTGCAATGAGAATTACC-3′

### Western blots

HEK293T cells were transfected with *EPHA4*-C1-pEGFP plasmid or the corresponding wild type vector to detect the missense variant in *EPHA4* (c.2546G>A, p.Cys849Tyr). Cells were cultured in six-well plates and transfected with DNA (2 mg/well) using Lipofectamine 3000 reagent (Thermo Fisher Scientific, Waltham, MA, USA). After 48 hours, cells were harvested, and protein extracts were prepared as described (Ding et al., 2017). The mutant proteins and wild-type proteins were fused to GFP. The expression of the two proteins was compared by western blots with an GFP antibody using the ECL detection system. The GFP antibody was purchased from Cell Signaling Technology (Cell Signaling, Danvers, MA, USA). The CDK5 antibody was purchased from Santa Cruz Biotechnology (Santa Cruz, CA, USA). A phospho-specific antibody against CDK5 phosphorylated at Tyr (GeneTex, Irvine, CA 92606 US) was purchased from GeneTex. Western blot experiments were repeated twice with similar results for the replicates.

### Genotyping and imputation of GWAS

In the East Asian Japanese cohort, genotyping was performed by Illumina Human610 Genotyping BeadChip, Illumina HumanOmniExpressExome and HumanOmniExpress as our previous GWASs (Kou et al., 2019; Kou et al., 2013; Ogura et al., 2015; Takahashi et al., 2011). For quality control (QC), subjects with call rate□<□0.98, high-degree relatedness with other subjects and outliers of East Asian ethnicity were excluded. For variants QC, the exclusion criteria of variants were as follows: call rate□<□0.99, P-value for Hardy-Weinberg equilibrium < 1.0□×□10^−6^ and minor allele count < 10. The reference panel for imputation, namely JEWEL7K, was composed of 1000 Genomes Project phase 3 (v5) (Auton et al., 2015) and Japanese whole-genome sequence data (Das et al., 2016) with 3256 high-depth subjects (≥30 read counts) and 4216 low-depth subjects (≤ 15 read counts). Using EAGLE 2.4.1 (https://alkesgroup.broadinstitute.org/Eagle/) to determine the haplotypes, pre-phasing was conducted. Genotypes were imputed using Minimac4 (version1.0.0) (Das et al., 2016). After imputation, we excluded variants with minor allele frequency (MAF) < 0.005 and low imputation quality (R^2^ < 0.3).

In the East Asian Hong Kong cohort, samples were genotyped with Illumina Human Omni ZhongHua-8 Beadchips. Illumina Genome Studio v2.0 was used to convert raw data into PLINK format. The QC steps of samples and variants were described in a previous study (Marees et al., 2018). Genotype phasing and imputing were executed using SHAPEIT v2.r900 (Delaneau et al., 2012) and IMPUTE2 (Marchini et al., 2007). The imputed data was filtered using the following parameters: INFO > 0.6, Certainty > 0.8, and MAF > 0.01. Association analysis was performed using PLINK v1.9 logistic regression with covariates: sex, age and top 20 principal components of the variance-standardized relationship matrix.

### East Asian GWAS meta-analysis for IS

For the meta-analysis of the four datasets (three Japanese datasets and one Hong Kong dataset), an inverse-variance-based method was performed by METAL (version2011-03-25) (Willer et al., 2010). SNPs in three or more of the four cohorts were used in subsequent analyses.

### Gene-based common variant analysis and eQTL analysis

SNPs in *EPHA4* or within 20 kb flanking *EPHA4* with a significant association with IS were retrieved. SNPs were matched to potential eQTLs according to the GTEx database (v8, https://gtexportal.org). Gene-based common variant analyses were performed using Multi-marker Analysis of GenoMic Annotation (MAGMA) (de Leeuw et al., 2015) and FUMA (Watanabe et al., 2017) using default settings with LD information from the 1000 Genomes Project East Asian population (1KGP EAS) as a reference. SNPs located within two kb upstream and one kb downstream from *EPHA4* were included in the gene-based analysis.

### Zebrafish strains, mutants, and morphants

Zebrafish Tuebingen (TU) strains were maintained at 28°C on a 14-hour/10-hour light/dark cycle. Embryos were raised at 28.5°C in E3 medium (5 mM NaCl, 0.17 mM KCl, 0.39 mM CaCl_2_, 0.67 mM MgSO_4_) following standard protocols. Zebrafish have two homologs of *EPHA4*, *epha4a* and *epha4b.* The CRISPR/Cas9 system was used to generate zebrafish *epha4a* and *epha4b* mutants. To increase efficiency, we injected Cas9 mRNA together with multiple single guide RNAs (sgRNAs) for each gene (Table S7). The sgRNA sequences for *epha4a* and *epha4b* are listed in Table S7. Morpholino sequences for *epha4a* and *efnb3b* knockdown analysis are also listed in Table S7.

### Micro CT imaging

Adult zebrafish *epha4a* mutants or wild-type siblings were euthanized with tricaine methanesulfonate and fixed in 4% paraformaldehyde. Micro CT images were captured using a PerkinElmer Quantum GX2. Planar images acquired over 360° of rotation were reconstructed using QuantumGX. Three-dimensional renders of the skeleton were made with Analyze 12.0 software (AnalyzeDirect).

### Behavioral recordings and analysis

For behavior analysis, individual zebrafish larvae were transferred into 24-well plate with fresh E3 medium at 8 dpf. Then, the plate was placed inside the Daniovision (Noldus) observation chamber for further behavior analysis. Video-tracking of swimming activity and further statistical analysis were performed using the EthoVision™ XT10 software. Behavioral data were shown as total swim distance (mm), average velocity (mm/s), average relative turn angle (degree), average absolute turn angle (degree) and average absolute angular velocity (degree/s) at a duration for 4 min. A positive angle reflects a leftward turn.

For tactile stimulation, zebrafish larvae were placed in a concave slide, and tactile stimulation was performed with glass capillaries to the head or tail sides. Larval startle responses were recorded using a high-speed video camera (Mikrotron, EoSens Mini1) at 1000 fps. Automated analysis of larval movement was performed using the FLOTE software package (Jain et al., 2014).

For optogenetic studies, the Tg(Gal4^s1020t^; *UAS*:ChR2) transgenic embryos were injected with control, *epha4a* or *efnb3b* morpholinos at one cell stage. At 5 dpf, the head of injected larva was mounted in 1% low melting-point agarose (Sigma) and the body was exposed in glass-bottom dishes (WPI). Leica M165FC fluorescence microscope was used to irradiate with a blue laser at 488nm wavelength. Larval responses were recorded using a high-speed video camera (Mikrotron, EoSens Mini1) at 500 fps. Body curvature analysis of larval movement was performed using the FLOTE software package.

### Whole-mount *in situ* hybridization and immunofluorescence

The primer sequences used to amplify *epha4a*, *epha4b* and *rfng* genes were listed in Table S7. Probe synthesis and whole-mount in situ hybridization were performed according to standard protocols. For reticulospinal neurons immunostaining, embryos were fixed in 2% trichloroacetic acid at 48 hpf for 3-4 h, washed twice in 0.5% Triton X-100 in PBS and blocked in 0.5% Triton X-100, 10% normal goat serum, 0.1% bovine serum albumin (Solarbio) in PBS for 1 h. The embryos were stained by monoclonal anti-neurofilament 160 antibody (Sigma-Aldrich) overnight at 4°C, then stained by goat anti-mouse Alexa Fluor 488 (Invitrogen) after washing.

### Analysis of neuronal calcium signals activity

Control, *epha4a* or *efnb3b* morpholinos were injected into Tg(*elavl3*:GAL4; UAS:GCaMP) transgenic embryos at one cell stage. At 24 hpf, embryos were paralyzed with 0.5 mg/ml α-Bungarotoxin (AlomoneLabs), then mounted in 1% low melting-point agarose (Sigma) in glass-bottom dishes (WPI). Neuronal calcium signal images were collected with IXON-L-888 EMCCD camera equipped on Dragonfly 200 Spinning Disk Confocal Microscope using a 20×/0.55 objective within 1 min at a frame rate of 10 fps.

Images were analyzed with imageJ software. To quantify the change in fluorescence intensity, a region of interest (ROI) was defined, and the fluorescence intensity F_t_ of different frames was normalized to F_t_/F_min_ based on the minimum fluorescence intensity F_min_ in all frames. To compare the left-right alternation pattern of neuronal calcium signals, the ratio of the left-side calcium signals frequency (f_left_) to the right-side calcium signals frequency (f_right_) was logarithmically transformed (lg-transformed).

### Quantification and statistical analysis

Statistical analyses were performed in SPSS (version 15.0). Unpaired Student’s t-test, Welch one-way ANOVA or two-way ANOVA followed by Turkey’s multiple comparison test were applied when appropriate. All experiments were replicated at least three times independently. P<0.05 was considered statistically significant.

### Study approval

Approval for the study was obtained from the ethics committee at the Peking Union Medical College Hospital (JS-098, JS-2364), the medical ethics committee of the Keio University Hospital (No. 20080129), the ethical committee of RIKEN Yokohama Institute (No. H20-17(8)), and the Institutional Review Board of the University Texas Southwestern Medical Center (protocol STU 112010L150). Written informed consent was obtained from each participating individuals and families in the three cohorts. For the control group, the protocols were approved by the ethics committee at Peking Union Medical College Hospital. All zebrafish studies were approved by the Animal Care Committee of the Ocean University of China (Animal protocol number: OUC2012316).

## Acknowledgments

We appreciate all the patients, their families and clinical research coordinators, including physicians who participated in this project. The authors also acknowledge the Texas Advanced Computing Center (TACC) at The University of Texas at Austin for providing computing resources that have contributed to the results related to the TSRHC cohort. URL: http://www.tacc.utexas.edu.

## Funding

National Key R&D Program of China 2023YFC2509700 (LW)

National Natural Science Foundation of China 81822030 (NW)

National Natural Science Foundation of China 82102522 (LW)

National Natural Science Foundation of China 31991194 & 32125015 (CZ)

National Natural Science Foundation of China 82172382 (TJZ)

CAMS Innovation Fund for Medical Sciences 2021-I2M-1-051 (TJZ, NW)

CAMS Innovation Fund for Medical Sciences 2021-I2M-1-052 (ZW)

CAMS Innovation Fund for Medical Sciences 2020-I2M-C&T-B-030 (TJZ)

Beijing Natural Science Foundation 7222133 (SW)

Non-profit Central Research Institute Fund of Chinese Academy of Medical Sciences No. 2019PT320025 (NW)

National High Level Hospital Clinical Research Funding 2022-PUMCH-D-004 (TJZ)

National High Level Hospital Clinical Research Funding 2022-PUMCH-C-033 (NW)

Shandong Natural Science Foundation ZR202102210113 (LW)

Shandong Province Taishan Scholar Project (LW)

Gabriella Miller Kids First Program grant X01 HL132375-01A1 (JR)

## Data and materials availability

The datasets used and/or analyzed during the current study are available from the corresponding author on reasonable request.

## Author contributions

Conceptualization: LW, SZ, CZ, NW

Methodology: LW, SZ, XY, PZ, WW, XC, AK, YK, NO, YL, CT, PL, CZ, NW

Investigation: LW, SZ, XY, PZ, WW, XC, AK, YK, JL, XF, NO, YL, LL, HX,

XL, SL, CT, SC, XZ, PL, SI, CZ, NW

Visualization: LW, SZ, XY, PZ, PZ, HX, JL, CZ, NW

Funding acquisition: LW, SW, TJZ, GQ, ZW, JR, CZ, NW

Project administration: LW, SZ, YN, ZW, GQ, TJZ, CZ, NW

Supervision: LW, SZ, SY, YN, ZW,GQ, TJZ, CZ, NW

Writing – original draft: LW, SZ, XY, JL, CZ, NW

Writing – review & editing: LW, SZ, XY, PZ, WW, KX, XC, QL, AK, YK, JL, XF, NO, YL, LL, HX, PZ, XL, YN, SW, SL, CT, ZL, SC, XZ, PL, JEP, ZW, GQ, SI, JRL, CAW, TJZ, CZ, NW

## Supplementary Materials

**Supplementary Fig. 1.**
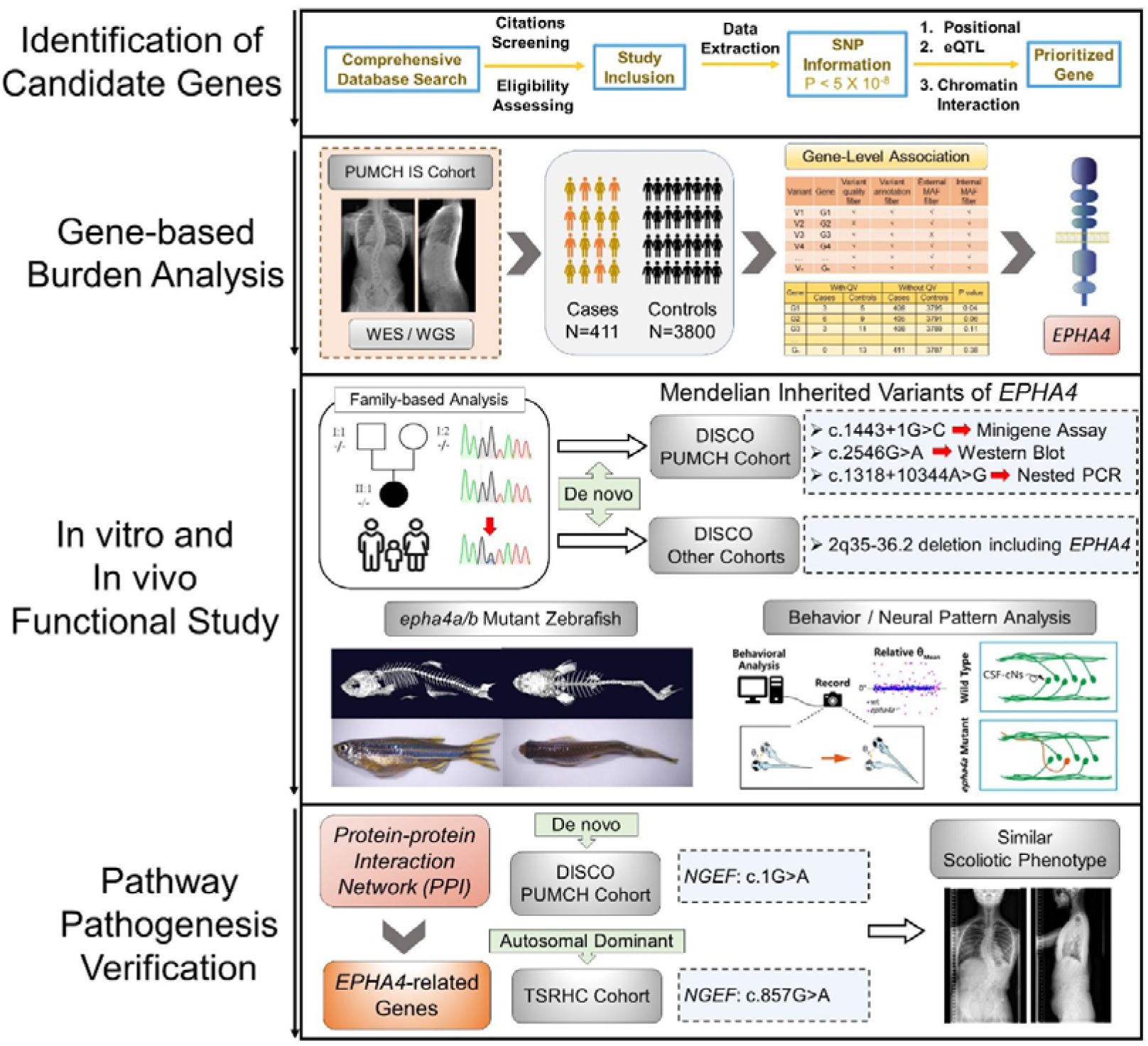
Flowchart for identification of causative genes. The processes of candidate gene mapping and gene-based burden analysis were described. The variants in *EPHA4* and *NGEF* identified in each cohort were displayed. Abbreviations: SNP (single nucleotide polymorphism); eQTL (expression quantitative trait locus); IS (idiopathic scoliosis).

**Supplementary Fig. 2.**
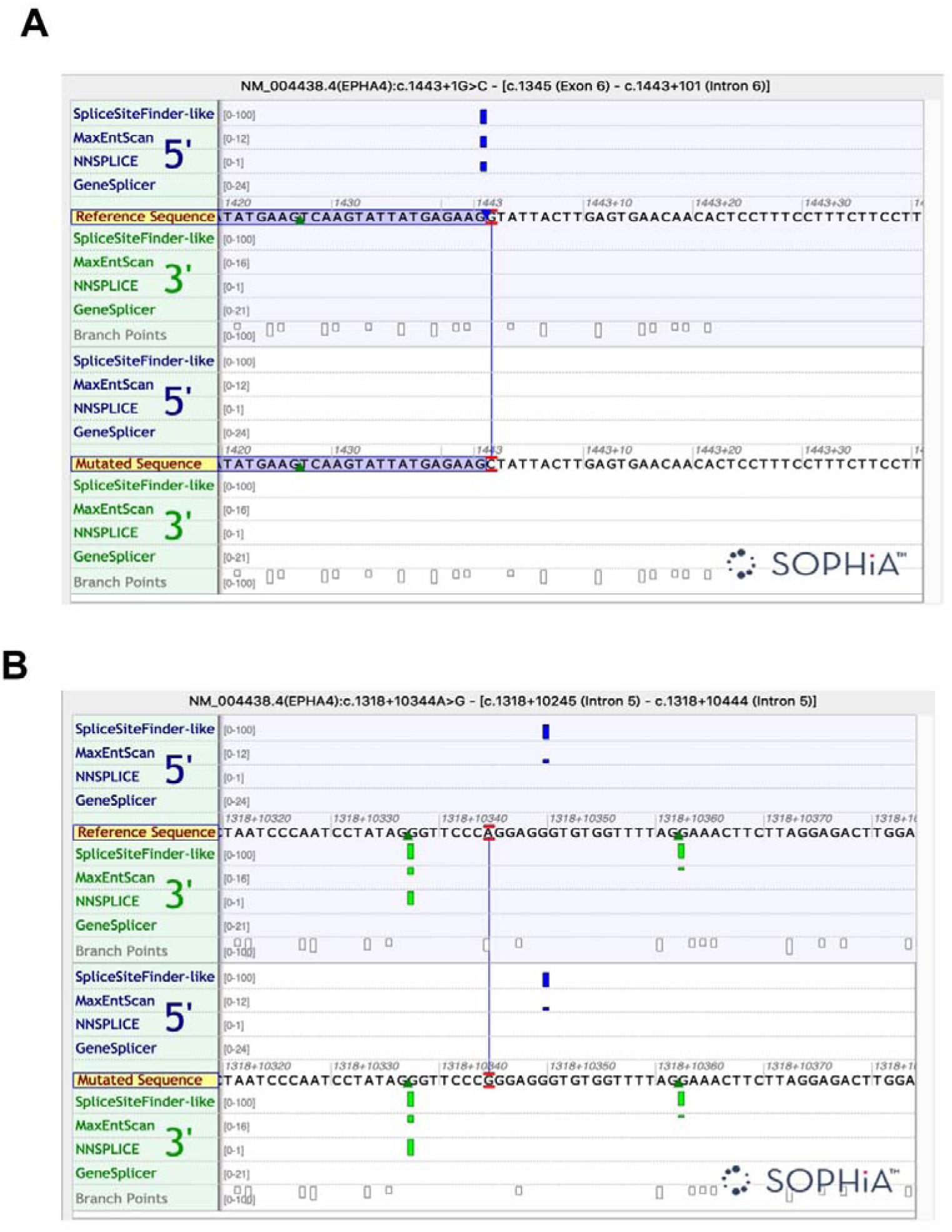
The splicing analysis by the Alamut software. The results of Alamut software by four algorithms (SpliceSiteFinder-like, MaxEntScan, NNSPLICE, and GeneSplicer).

**Supplementary Fig. 3.**
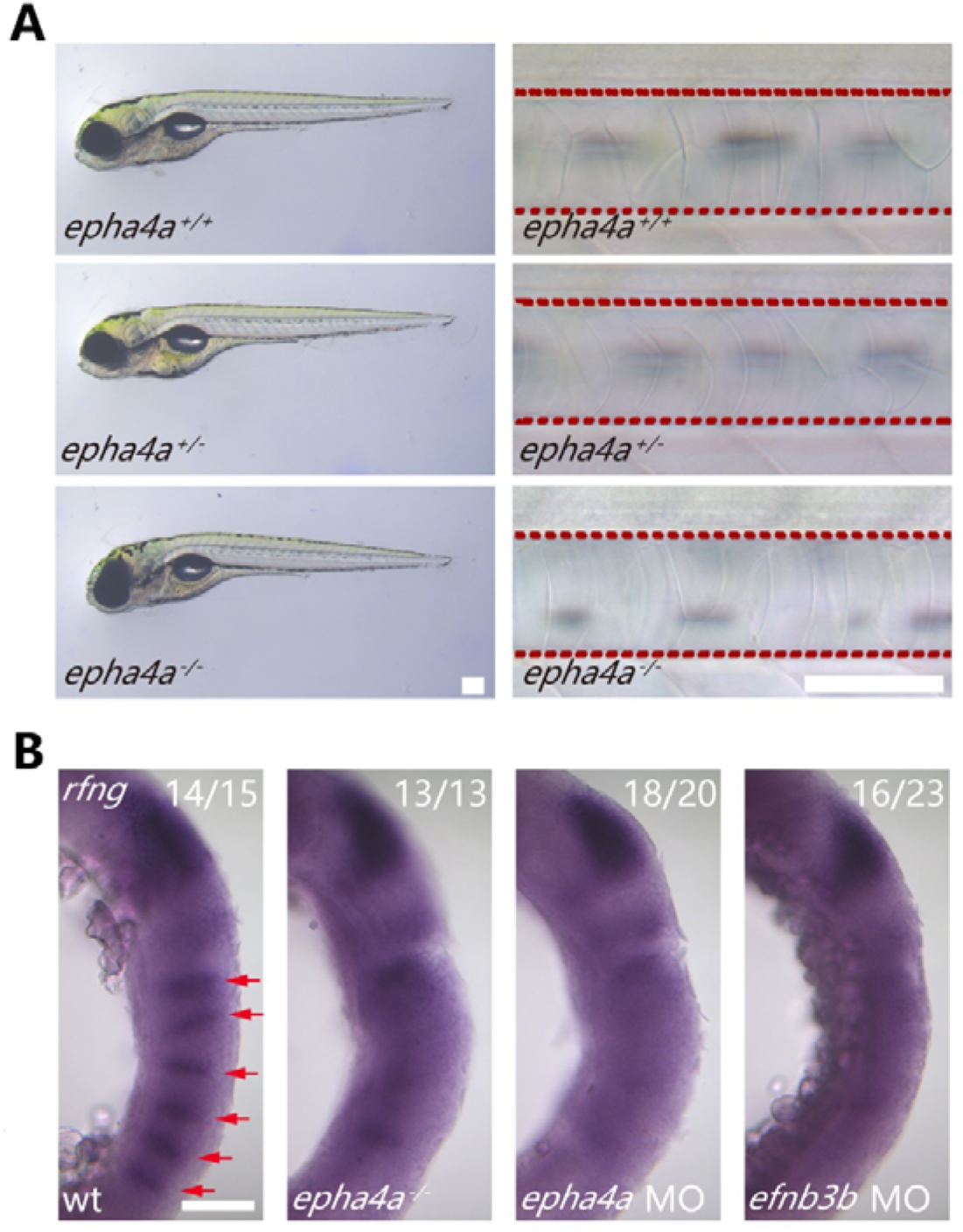
Phenotypes of *epha4a* mutants. **A:** Representative images showing the notochord and external phontypes of wild-type and *epha4a* mutants. The red dashed lines indicate the area of the notochord. **B:** Whole-mount in situ hybridization results showing the expression of *rfng* gene in 18 hpf wild type, homozygous mutant or morphant larvae as indicated. *rfng* gene marks boundary cells in the hindbrain. The red arrows indicate boundary expression. The numbers of embryos analyzed are shown on top right. Scale bars: 200 µm in panel (A), 100 µm in panel (B).

**Supplementary Fig. 4.**
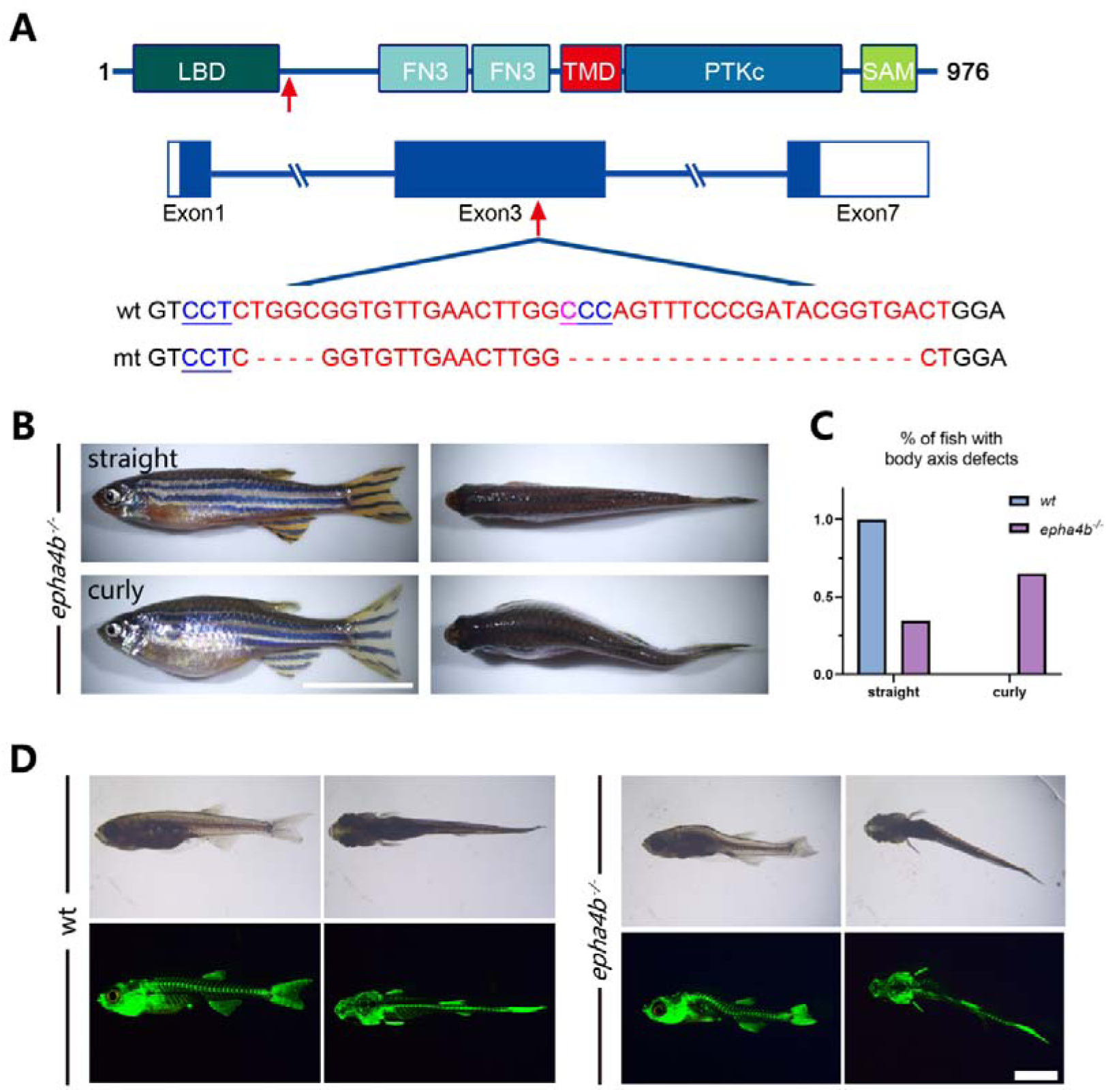
Zebrafish *epha4b* mutants exhibited body axis defects during development. **A:** Diagram showing the protein domains, genomic structures, and sequences of the wild-type and corresponding *epha4b* mutants. Red arrows indicate mutation sites. **B:** Representative images of *epha4b* mutants. **C:** Bar graph showing the percentages of adult fish with normal and body axis defects in wild-type (n=39) and *epha4b* mutants (n=43). **D:** Bright-field and GFP fluorescent images showing scoliosis *epha4b* mutants at 28 dpf as indicated by Tg*(Ola.Sp7:NLS-GFP)* transgene, which labels the bone skeleton. Scale bars: 1 cm in panel (B), 2 mm in panel (D).

**Supplementary Fig. 5.**
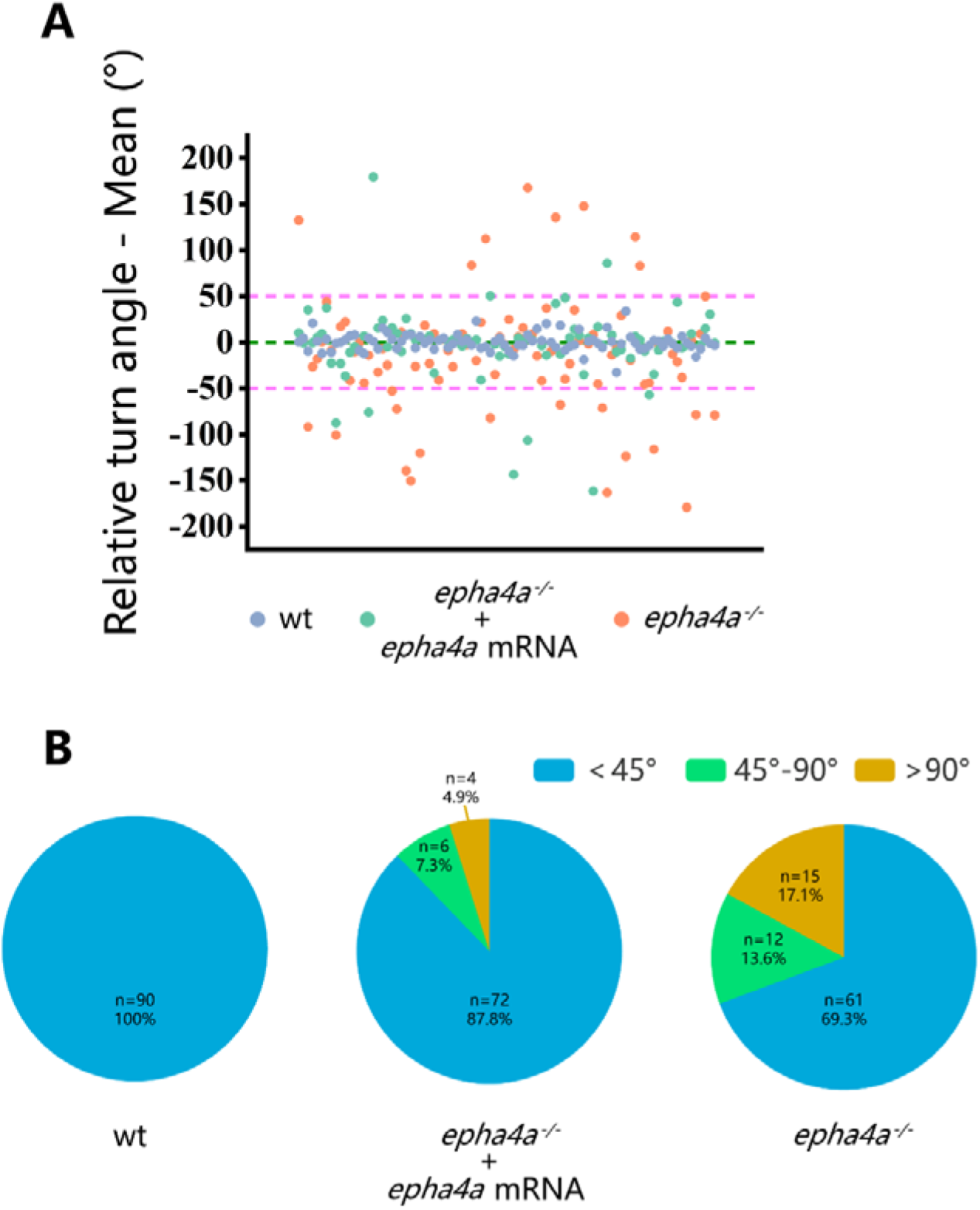
Rescue of the *epha4a* homozygous mutant phenotype by *epha4a* mRNA injection. A: Scatter plot showing the relative turning angle of wild-type, *epha4a* homozygous larvae and homozygous larvae injected with the corresponding mRNA. B: The pie chart shows the proportional distribution of relative turning angles in three groups of larvae. N=30 larvae, n=90 experiments. Larvae that remained stationary during the recording were excluded.

**Supplementary Fig. 6.**
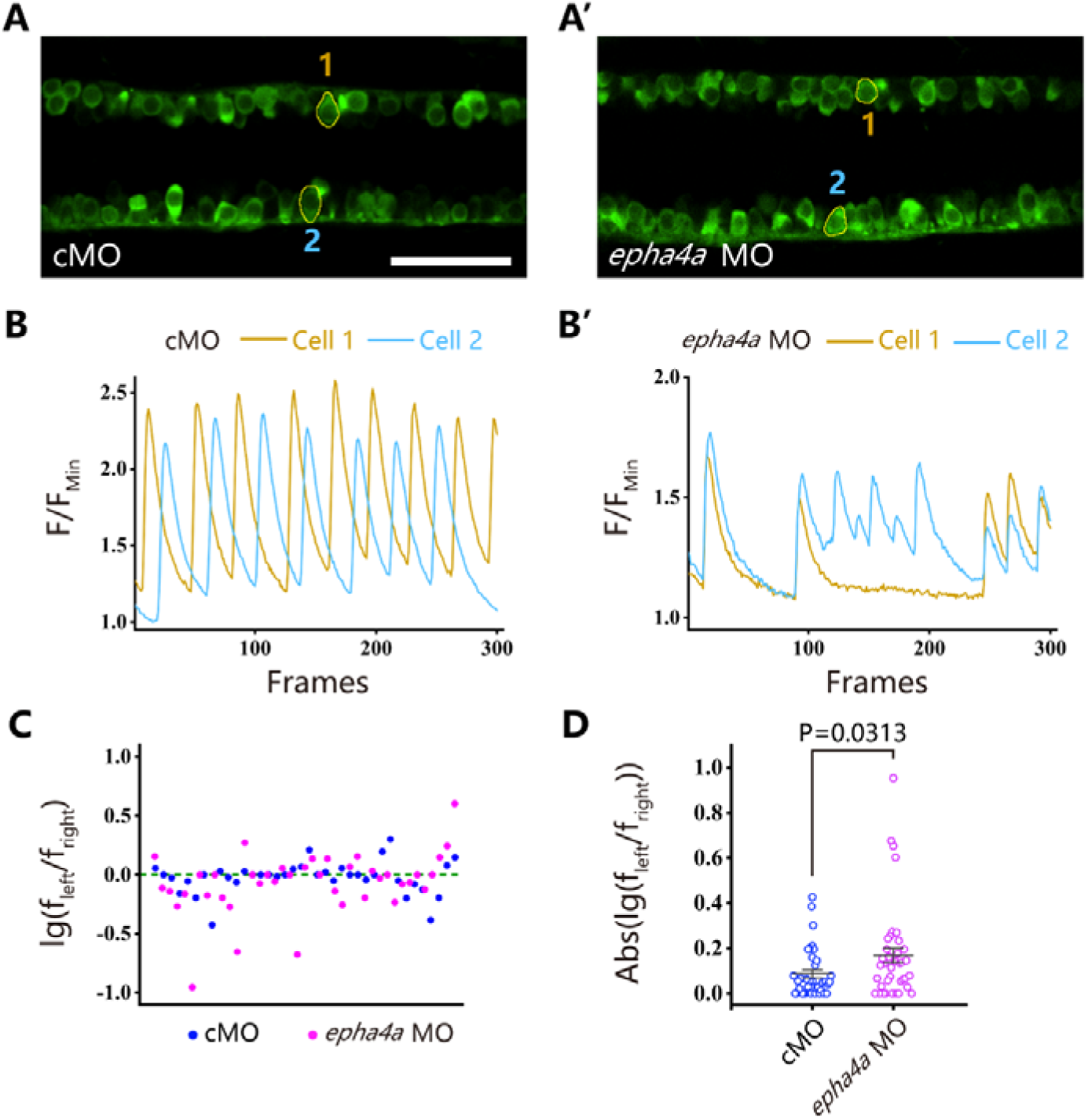
Uncoordinated left-right activation of spinal cord neurons in the absence of Epha4a. A, A’: Fluorescent images showing the dorsal-view of 24 hpf Tg(*elavl3*:GAL4; *UAS*:GCaMP) double transgenic larvae. The corresponding movies are shown in Movie S7 and S8. **B, B’:** Line charts showing the quantification of fluorescence changes of the region of interests (ROIs, circled in A, A’) in control and *epha4a* morphants. **C:** Scatter plot showing the distribution trend of the ratio of the calcium signal frequency between left and right in control (N=15 larvae, n=38 experiments) and *epha4a* morphants (N=15 larvae, n=41 experiments). **D:** Statistical graph of the ratio of the calcium signal frequency between left and right in control and *epha4a* morphants. Scale bars: 50 μm in panel **(A, A’)**.

**Supplementary Fig. 7.**
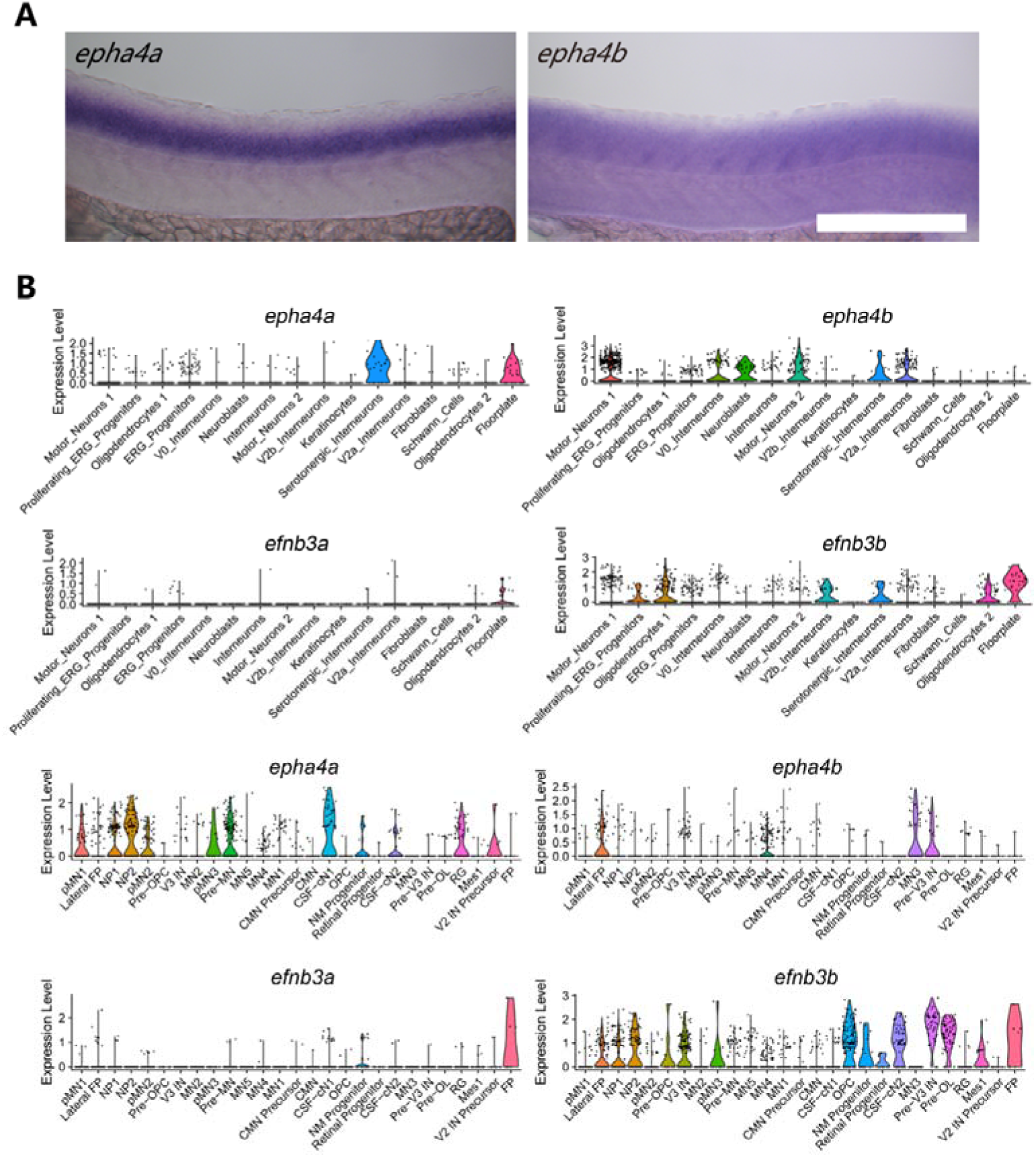
Expression pattern of *epha4* and *efnb3* in the spinal cord. **A:** Whole-mount in situ hybridization results showing the expression of *epha4a* and *epha4b* genes in the spinal cord in 24 dpf zebrafish larvae. **B:** Violin plot showing *epha4a, epha4b, efnb3a,* and *efnb3b* gene expression in different cell types. Single cell transcriptome data were acquired from two different studies (top and bottom) (Cavone et al., 2021; Scott et al., 2021). Scale bars: 200 µm in panel (A).

**Supplementary Fig. 8.**
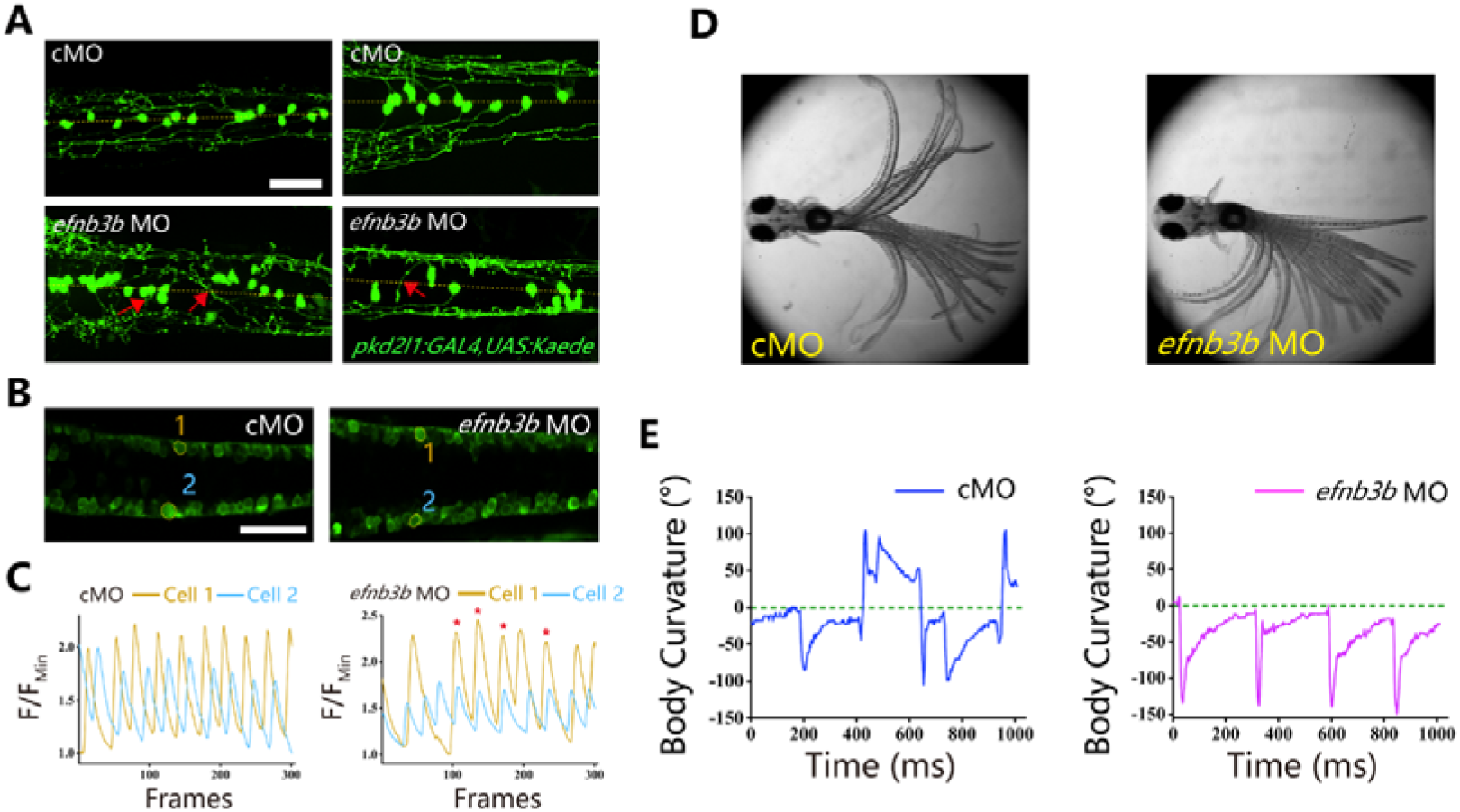
**Left-right coordination defects in *efnb3b* morphants**. **A:** Fluorescent images showing the distribution of ascending axons of CSF-cNs marked by Tg*(pkd2l1:GAL4,UAS:Kaede)* in 2 dpf *efnb3b* morphants. The red arrows indicate aberrantly extended axons in *efnb3b* morphants. **B:** Fluorescent images showing the dorsal-view of 24 hpf Tg(*elavl3*:GAL4; *UAS*:GCaMP) transgenic larvae. **C:** The line chart showing the quantification of fluorescence changes of the ROIs in control morphants and *efnb3b* morphants as indicated in panel B. **D:** Superimposed frames of tail oscillations in 5dpf control and *efnb3b* morphants. **E:** A plot of body curvature angles in panel (D). The positive angle means turning right. Scale bars: 50 μm in panel (A, B).

**Supplementary Fig. 9.**
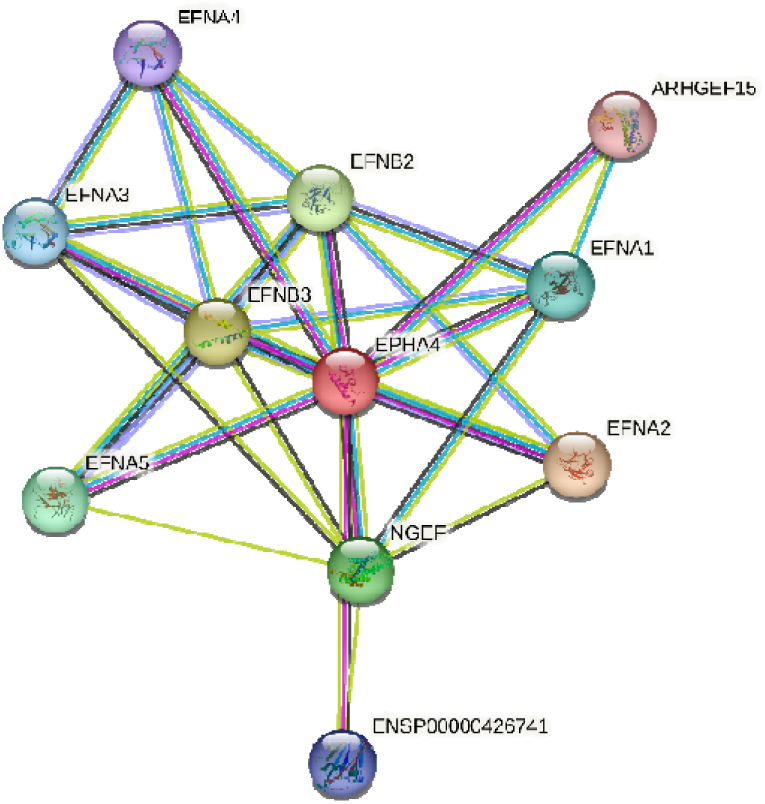
Proteins in STRING that interact with EPHA4. Protein interaction network analysis, as illustrated by STRING v11.5, was used to identify rare variants in genes related to *EPHA4*. Known and predicted protein-protein interactions are included in the STRING database. Multiple proteins, including Ephexin, are associated with EPHA4 physically or functionally. Ephexin, encoded by *NGEF*, is a neuronal guanine exchange factor.

**Supplementary Table 1.**
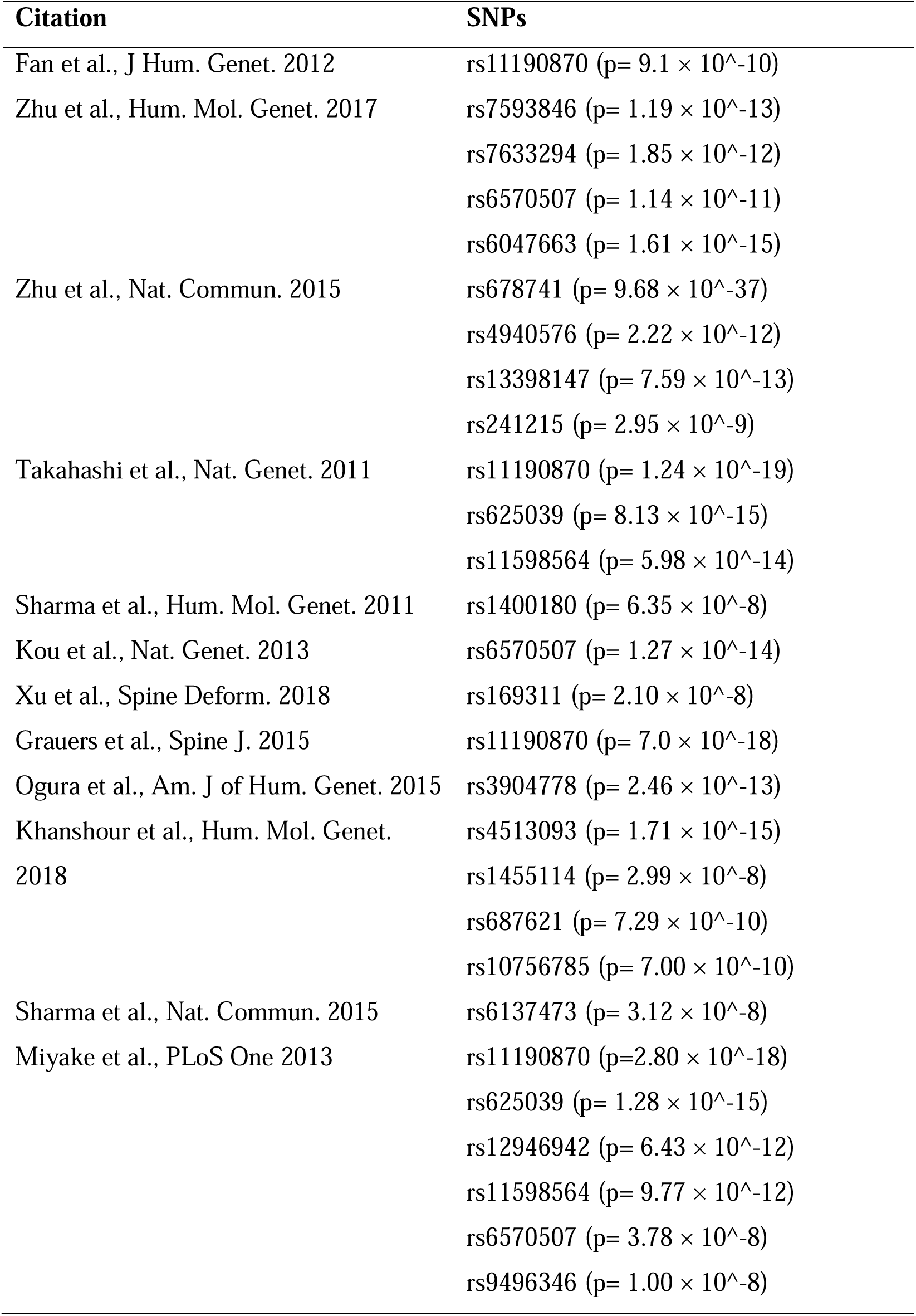
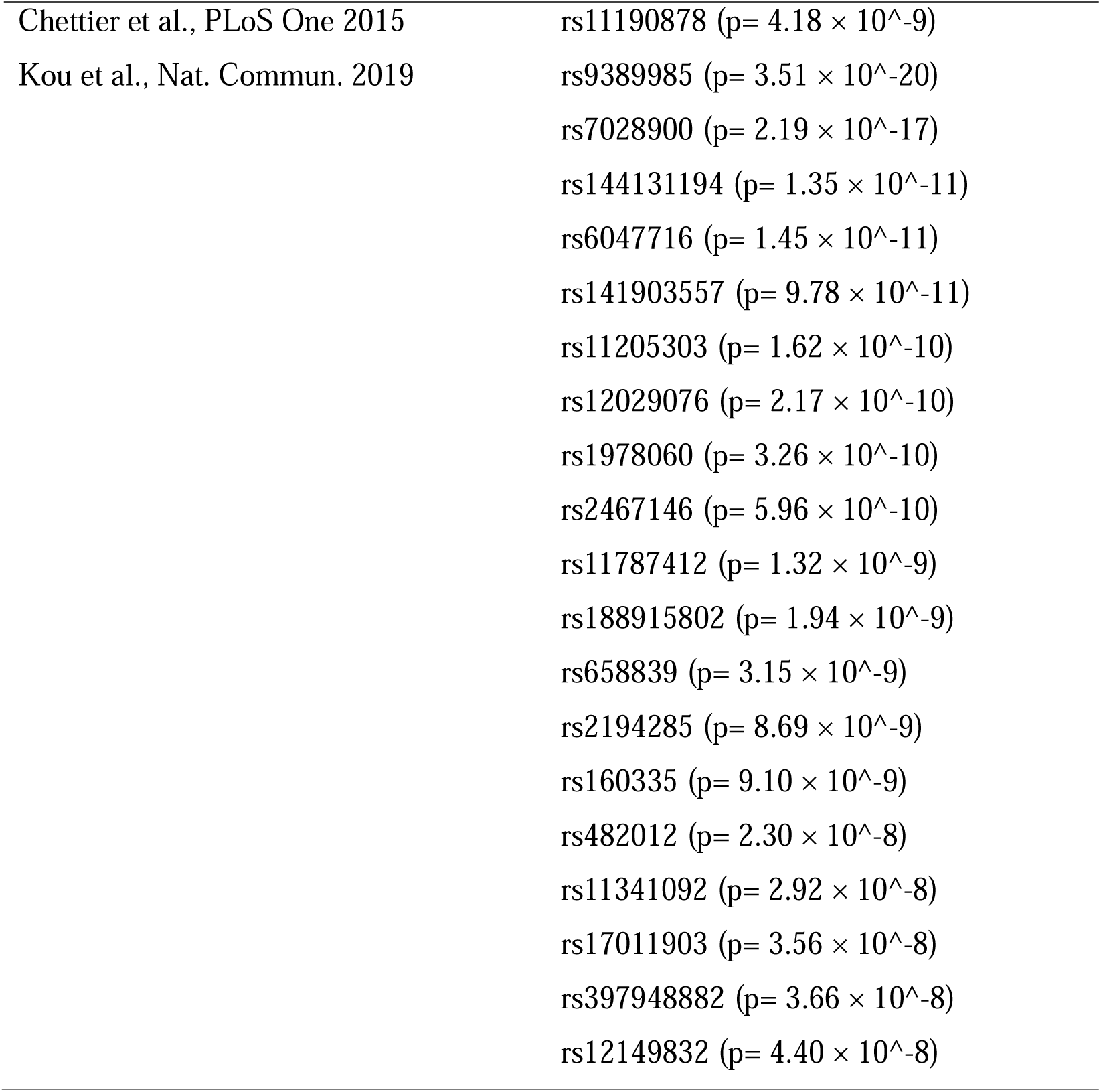
Summary of the 14 studies and their corresponding SNPs included in the candidate genes mapping.

| Citation | SNPs |
| --- | --- |
| Fan et al., J Hum. Genet. 2012 | rs11190870 ($p= 9.1 \times 10^{-10}$) |
| Zhu et al., Hum. Mol. Genet. 2017 | rs7593846 ($p= 1.19 \times 10^{-13}$) |
| | rs7633294 ($p= 1.85 \times 10^{-12}$) |
| | rs6570507 ($p= 1.14 \times 10^{-11}$) |
| | rs6047663 ($p= 1.61 \times 10^{-15}$) |
| Zhu et al., Nat. Commun. 2015 | rs678741 ($p= 9.68 \times 10^{-37}$) |
| | rs4940576 ($p= 2.22 \times 10^{-12}$) |
| | rs13398147 ($p= 7.59 \times 10^{-13}$) |
| | rs241215 ($p= 2.95 \times 10^{-9}$) |
| Takahashi et al., Nat. Genet. 2011 | rs11190870 ($p= 1.24 \times 10^{-19}$) |
| | rs625039 ($p= 8.13 \times 10^{-15}$) |
| | rs11598564 ($p= 5.98 \times 10^{-14}$) |
| Sharma et al., Hum. Mol. Genet. 2011 | rs1400180 ($p= 6.35 \times 10^{-8}$) |
| Kou et al., Nat. Genet. 2013 | rs6570507 ($p= 1.27 \times 10^{-14}$) |
| Xu et al., Spine Deform. 2018 | rs169311 ($p= 2.10 \times 10^{-8}$) |
| Grauers et al., Spine J. 2015 | rs11190870 ($p= 7.0 \times 10^{-18}$) |
| Ogura et al., Am. J of Hum. Genet. 2015 | rs3904778 ($p= 2.46 \times 10^{-13}$) |
| Khanshour et al., Hum. Mol. Genet. 2018 | rs4513093 ($p= 1.71 \times 10^{-15}$) |
| | rs1455114 ($p= 2.99 \times 10^{-8}$) |
| | rs687621 ($p= 7.29 \times 10^{-10}$) |
| | rs10756785 ($p= 7.00 \times 10^{-10}$) |
| Sharma et al., Nat. Commun. 2015 | rs6137473 ($p= 3.12 \times 10^{-8}$) |
| Miyake et al., PLoS One 2013 | rs11190870 ($p= 2.80 \times 10^{-18}$) |
| | rs625039 ($p= 1.28 \times 10^{-15}$) |
| | rs12946942 ($p= 6.43 \times 10^{-12}$) |
| | rs11598564 ($p= 9.77 \times 10^{-12}$) |
| | rs6570507 ($p= 3.78 \times 10^{-8}$) |
| | rs9496346 ($p= 1.00 \times 10^{-8}$) |
| Chettier et al., PLoS One 2015 | rs11190878 ($p= 4.18 \times 10^{-9}$) |
| Kou et al., Nat. Commun. 2019 | rs9389985 ($p= 3.51 \times 10^{-20}$) |
| | rs7028900 ($p= 2.19 \times 10^{-17}$) |
| | rs144131194 ($p= 1.35 \times 10^{-11}$) |
| | rs6047716 ($p= 1.45 \times 10^{-11}$) |
| | rs141903557 ($p= 9.78 \times 10^{-11}$) |
| | rs11205303 ($p= 1.62 \times 10^{-10}$) |
| | rs12029076 ($p= 2.17 \times 10^{-10}$) |
| | rs1978060 ($p= 3.26 \times 10^{-10}$) |
| | rs2467146 ($p= 5.96 \times 10^{-10}$) |
| | rs11787412 ($p= 1.32 \times 10^{-9}$) |
| | rs188915802 ($p= 1.94 \times 10^{-9}$) |
| | rs658839 ($p= 3.15 \times 10^{-9}$) |
| | rs2194285 ($p= 8.69 \times 10^{-9}$) |
| | rs160335 ($p= 9.10 \times 10^{-9}$) |
| | rs482012 ($p= 2.30 \times 10^{-8}$) |
| | rs11341092 ($p= 2.92 \times 10^{-8}$) |
| | rs17011903 ($p= 3.56 \times 10^{-8}$) |
| | rs397948882 ($p= 3.66 \times 10^{-8}$) |
| | rs12149832 ($p= 4.40 \times 10^{-8}$) |

**Supplementary Table 2.**
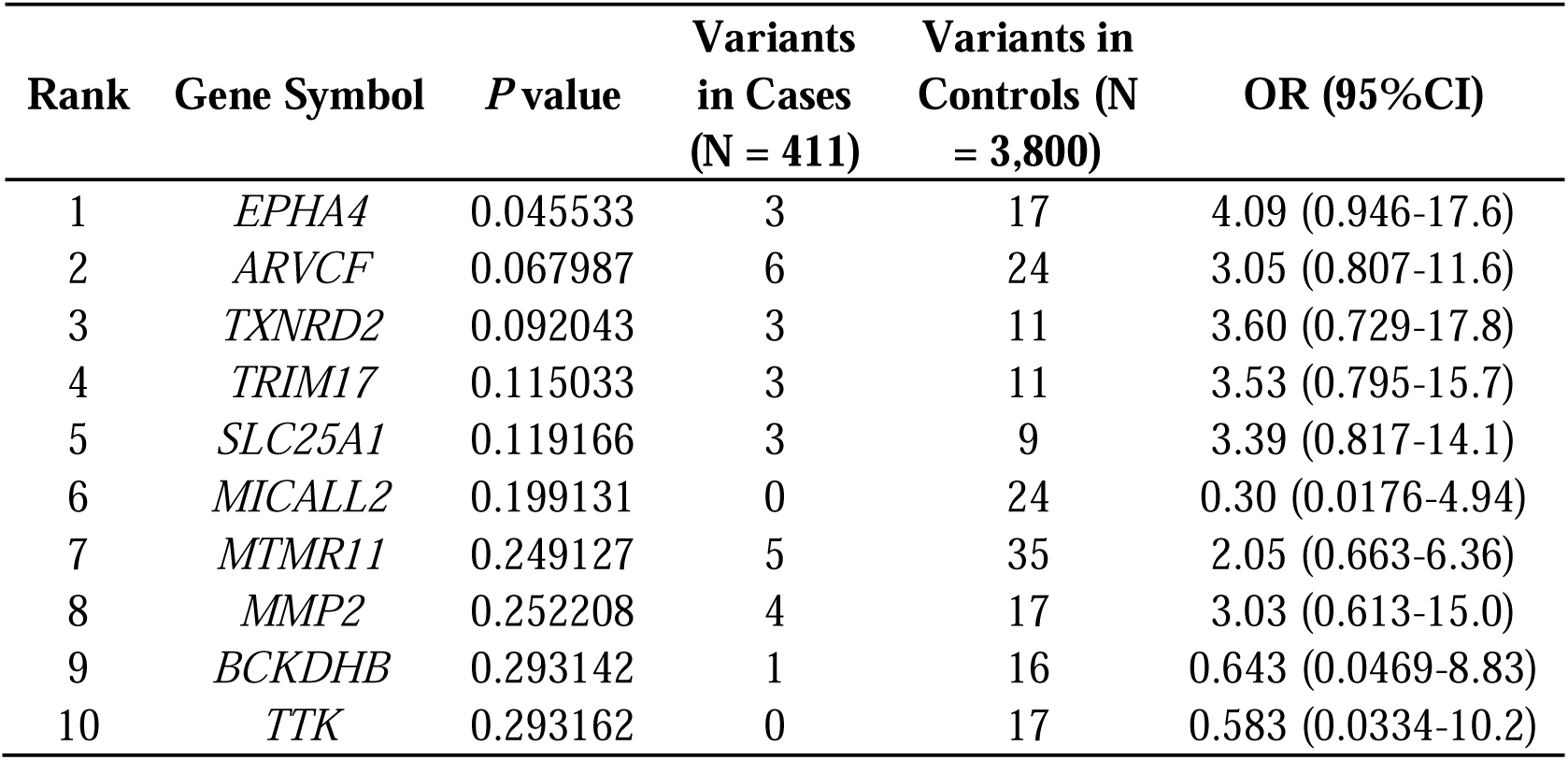
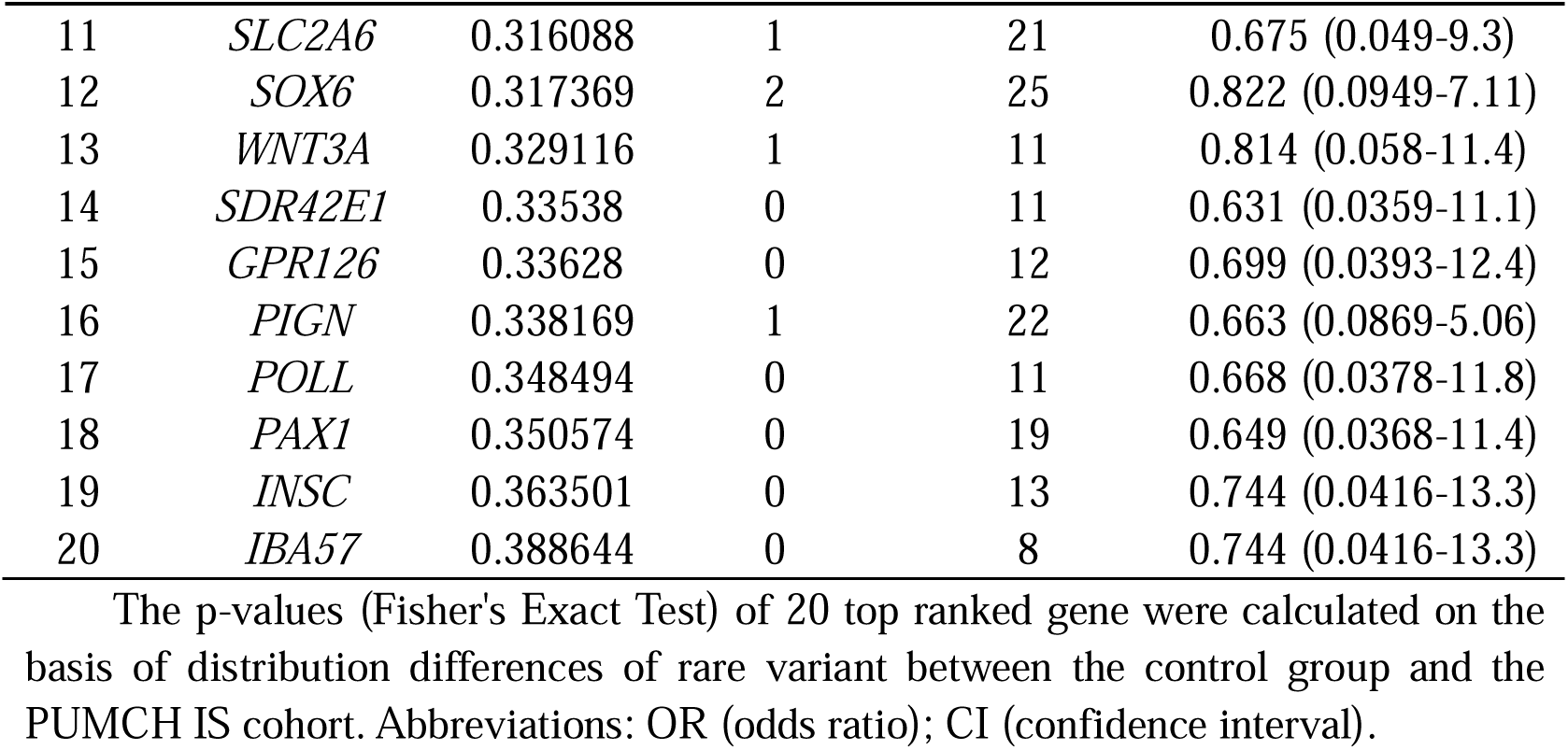
Summarized results of burden analysis.

**Supplementary Table 4.**
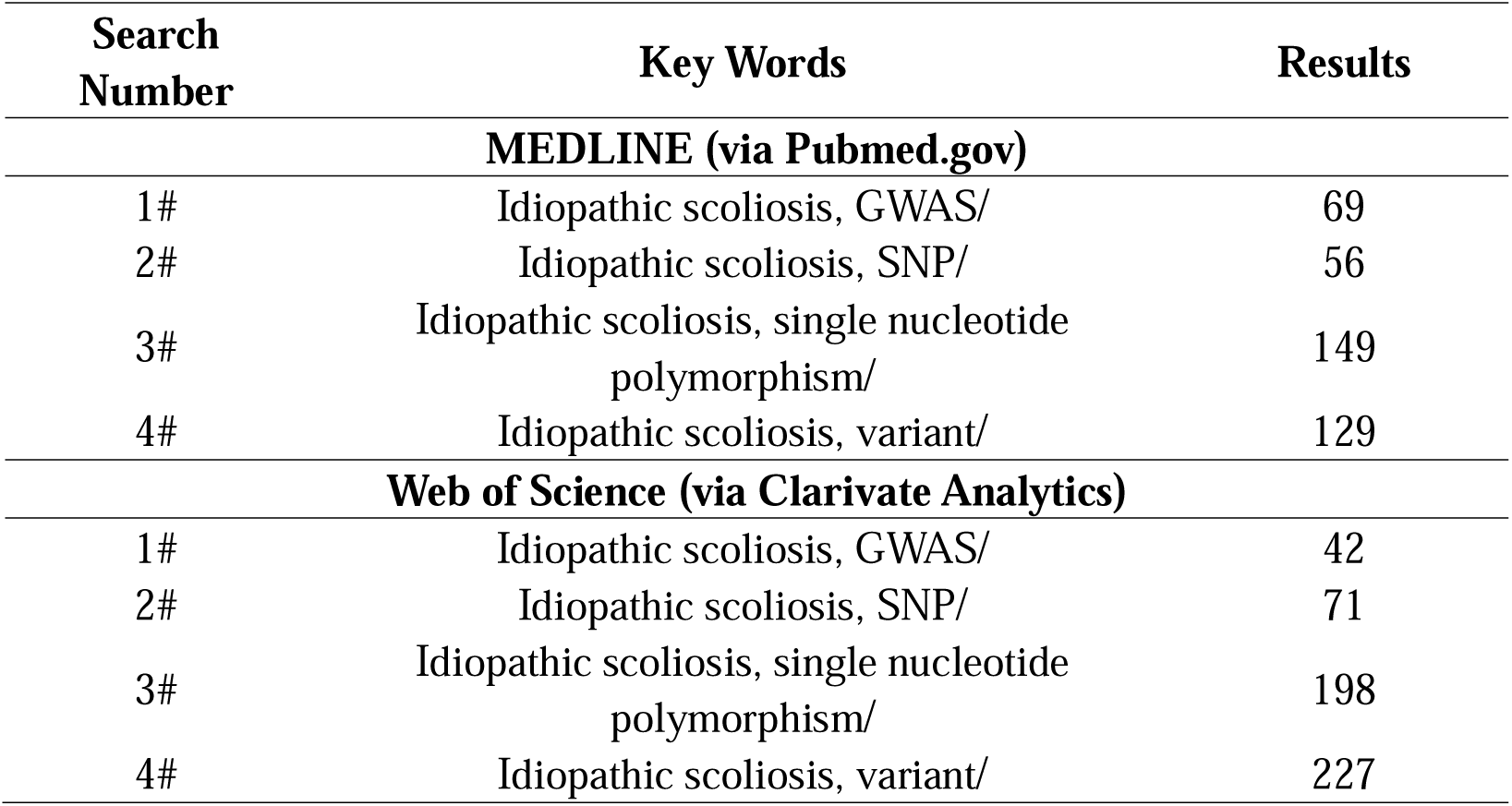
Search strategies for each database.

**Supplementary Table 5.** Inclusion and exclusion criteria of literature review.

|  | **Inclusion Criteria** | **Exclusion Criteria** |
| --- | --- | --- |
| Patient | Idiopathic scoliosis | Congenital scoliosis |
| Recruitment | Adolescent idiopathic scoliosis | Neuromuscular scoliosis |
|  | Early-onset idiopathic scoliosis | Syndromic scoliosis |
|  |  | Secondary scoliosis |
|  |  | Degenerative scoliosis |
| Study Type | Genome Wide Association | Linkage study |
|  | Study (GWAS) | Candidate gene study |
|  | Meta-analysis of GWAS | Exome sequencing |
|  |  | Whole genome sequencing |
| Study Design | Case-control | - |

**Supplementary Table 6.** Sequencing information of PUMCH IS cohort.

| **Characteristics** | **Cases** |  | **Controls** |  |
| --- | --- | --- | --- | --- |
|  | **Trio** | **Singleton** | **Trio** | **Singleton** |
| Exome sequencing | 41 | 113 | 2,021 | 224 |
| Genome sequencing | 116 | 148 | 483 | 1,072 |
| Sum | 157 | 261 | 2,504 | 1,296 |
| Passing QC | 155 | 256 | 2,504 | 1,296 |

Abbreviations: QC (quality control); IS (idiopathic scoliosis).

**Supplementary Table 7.**
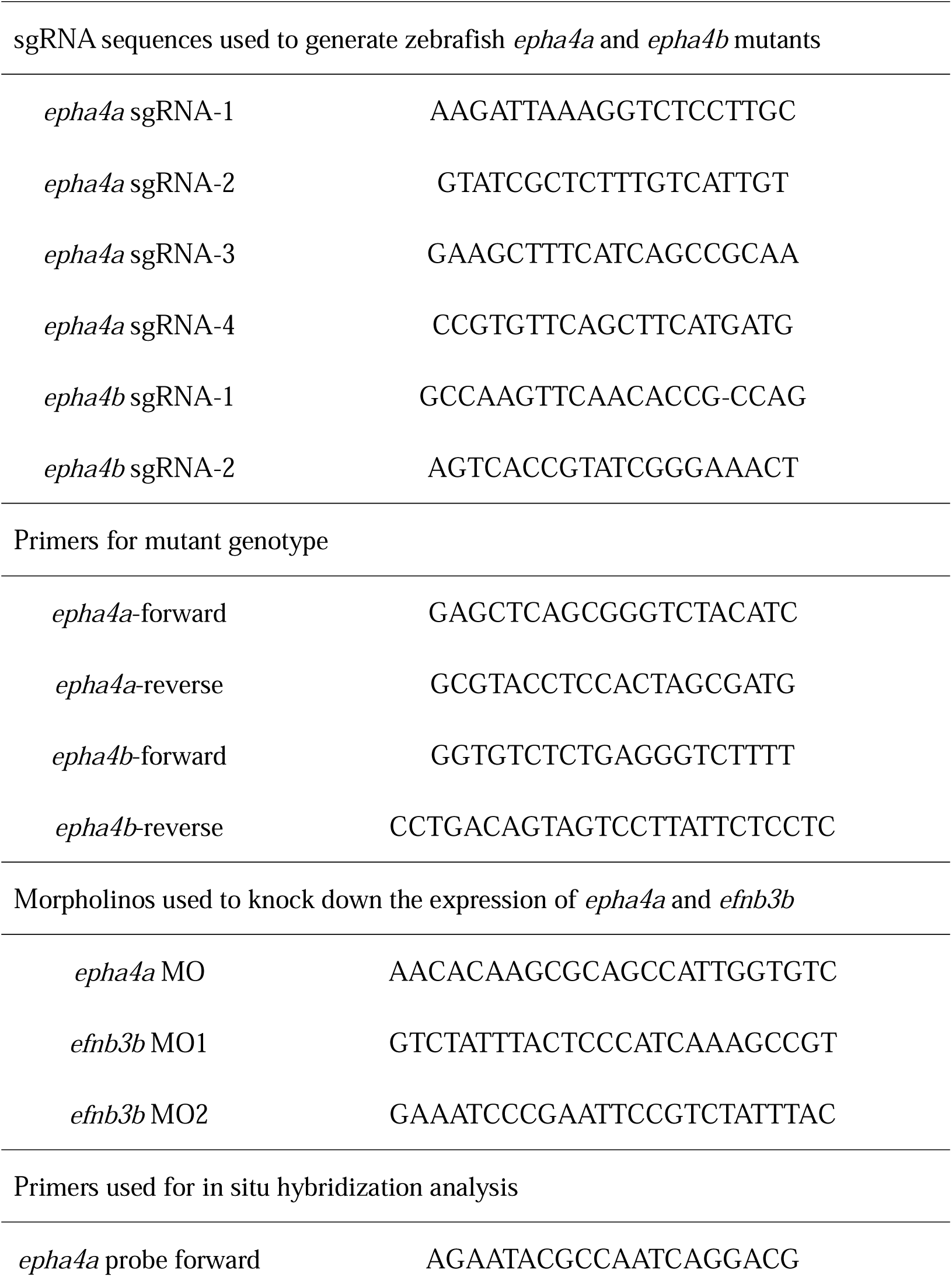
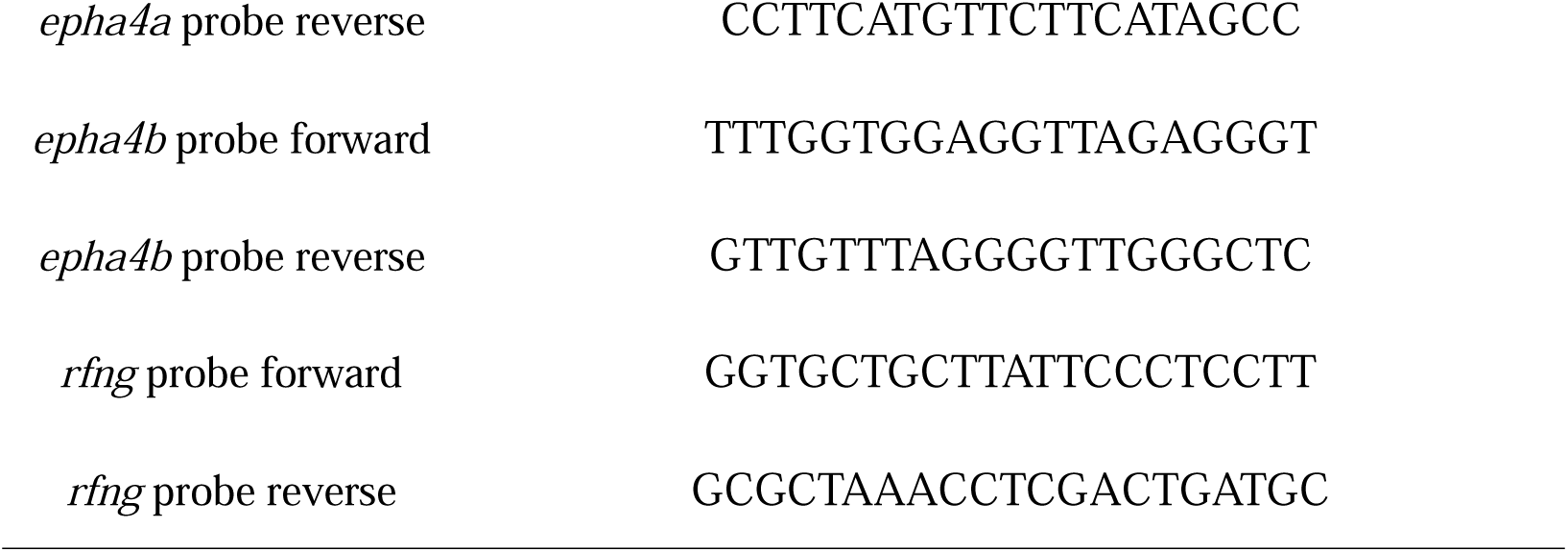
Sequences information of sgRNA and primers for zebrafish study.

**Supplementary Movie 1:** Video showing the swimming of three wild-type fish.

**Supplementary Movie 2:** Video showing the swimming of three *epha4a* mutants with severe or mild scoliosis.

**Supplementary Movie 3:** High-speed video showing the startle response in a wild-type larva at 5 dpf triggered by head tactile stimulation. Time units: ms.

**Supplementary Movie 4:** High-speed video showing the abnormal startle response of an *epha4a* mutant larva at 5 dpf triggered by head tactile stimulation. Time units: ms.

**Supplementary Movie 5:** Alternated activation of calcium signaling in motor neurons of 24 hpf wild type Tg(*elavl3*:GAL4; *UAS*: GCaMP) transgenic larva.

**Supplementary Movie 6:** Alternated activation of calcium signaling in motor neurons of 24 hpf wild type Tg(*elavl3*:GAL4; *UAS*: GCaMP) transgenic larva injected with control MO.

**Supplementary Movie 7:** Abnormal activation of calcium signaling in motor neurons of 24 hpf Tg(*elavl3*:GAL4; *UAS*: GCaMP) transgenic larva with homozygous *epha4a* mutant background.

**Supplementary Movie 8:** Abnormal activation of calcium signaling in motor neurons of 24 hpf Tg(*elavl3*:GAL4; *UAS*: GCaMP) transgenic larva injected with *epha4a* MO.

**Supplementary Movie 9:** Tail oscillation after light activation of 5dpf Tg(Gal4^s1020t^; *UAS*:ChR2) double transgene larva.

**Supplementary Movie 10:** Tail oscillation after light activation of 5dpf Tg(Gal4^s1020t^; *UAS*:ChR2) double transgene larva injected with *epha4a* MO.

